# The *Arabidopsis thaliana* TRAPPIII subunit AtTRAPPC8/AtTRS85 is involved in ER functioning and autophagy

**DOI:** 10.1101/2024.01.11.575191

**Authors:** Marta Hoffman-Sommer, Natalia Piłka, Anna Anielska-Mazur, Julita Nowakowska, Yasin Dagdas, Ewa Swiezewska

## Abstract

TRAPP (*transport protein particle*) tethering complexes are known for their function as Rab-GTPase exchange factors (GEFs). Two versions of the complex are considered functionally separate: TRAPPII, an activator of GTPases of the Rab11 family (RabA in plants) which functions in post-Golgi sorting, and TRAPPIII, activating the Rab1 family (RabD in plants) which regulates ER-to-Golgi trafficking and autophagy. In *Arabidopsis thaliana*, the TRAPPIII complex has been identified and its subunit composition established, but little is known about its functions. Here, we found that binary subunit interactions of the plant TRAPPIII complex are analogous to those of metazoan TRAPPIII, with the two large subunits TRAPPC8 and -C11 linking the TRAPP core and the small C12-C13 dimer. To gain insight into the functions of TRAPPIII in plants, we characterized two *A. thaliana trappc8* mutants. The mutants display abnormalities in plant morphology, in particular in flower and seed development. They also have autophagic defects, constitutive ER stress response, and elevated levels of the ER lipid dolichol – an indispensable cofactor of protein glycosylation. These results show that plant TRAPPC8 is involved in multiple trafficking steps in the cells and they suggest a novel link between ER membrane turnover and dolichol levels.

**HIGHLIGHT:** *Arabidopsis thaliana* TRAPPC8 is necessary for correct functioning of the ER, in particular for its lipid homeostasis. Dysruption of *TRAPPC8* leads to defects in secretion, autophagosome formation, and plant development.

## INTRODUCTION

Intracellular membrane traffic is critically important for the correct functioning of all eukaryotic cells because it maintains organelle identity. The existence of separate compartments in a cell is dependent on the tight regulation of transport events which must ensure the delivery of various compounds from the sites of their synthesis or entry into the cell to precisely defined destination sites (the sites of their activity and/or deposition). Rab GTPases play a crucial role in these processes: they contribute to membrane identity and they provide specificity by recruiting appropriate effector proteins (Elliott et al., 2020; Minamino & Ueda, 2019; Nielsen, 2020; Pfeffer, 2013). Localization of a specific Rab to the correct membrane and its activation is dependent on a broad set of regulatory proteins, among which are guanine-nucleotide exchange factors (GEFs) (Lamber et al., 2019) which activate their cognate Rabs by catalyzing the exchange of a Rab-bound GDP molecule for a fresh GTP molecule.

The TRAnsport Protein Particle (TRAPP) complexes TRAPPII and -III both function as GEFs for the Ypt1/Rab1 or/and Ypt31/32/Rab11 families. They differ from each other by subcellular localization and by substrate specificity towards members of the two Rab families. In yeast TRAPPIII activates Ypt1 (Thomas et al., 2018) while TRAPPII activates Ypt31/32 (Thomas & Fromme, 2016); in animal cells TRAPPII likely activates members of both Rab families while TRAPPIII is Rab1-specific (Riedel et al., 2018; Harris et al., 2021). The TRAPP complex subunits are conserved in eukaryotes from yeast to animals and plants, though higher eukaryotes and some fungi have additional subunits in TRAPPIII, absent in yeast cells (TRAPPC11, -12, -13) (Scrivens et al., 2011; Choi et al., 2011; Bassik et al., 2013; Pinar et al., 2019), and one TRAPPIII subunit (TRAPPC8) has only partial similarity to its yeast counterpart, Trs85p. Further, a plant-specific component has been identified in TRAPPII (TRIPP; Garcia et al., 2020), showing that the plant and animal complexes also differ from each other.

In this work we concentrated on TRAPPIII. The structures of yeast and metazoan TRAPPIII have been solved (Tan et al., 2013; Galindo et al., 2021; Joiner et al., 2021), so the positioning of the additional metazoan subunits C8-11-12-13 in relation to the common TRAPP core is known. In *Drosophila melanogaster*, the large subunits C8 and C11 interact with each other on one end and attach to the TRAPP core on their opposing ends (with C8 connecting to the core through the subunit C2, and C11 through C2L), while the smaller subunits C12-C13 form a heterodimer that attaches to the C8-C11 joint. Rab1-binding by the TRAPP core is thought to be facilitated by the conserved N-terminal arm of the C8 subunit (Galindo et al., 2021).

Plant TRAPPs are not as well described as the yeast and metazoan complexes. Their subunit composition has been found to be similar to that of the metazoan complexes (with one additional plant-specific subunit in TRAPPII) (Kalde et al., 2019; Garcia et al., 2020), but no structural data has been collected yet and their substrate specificity also remains uncertain. Currently available data suggest that TRAPPII activates not only members of the RabA (homologous to the animal Rab11 family) and RabD (homologous to the Rab1 family) clades, but also members of the B and E clades (Kalde et al., 2019). Data obtained previously for the C11 subunit of TRAPPIII suggest a possible role as a membrane tether at the TGN/EE (Rosquete et al., 2019 a, b), possibly as a GEF for RabD2. In this scenario, TRAPPIII would be involved in Golgi and post-Golgi trafficking, and endocytic sorting and recycling of specific components. Another function was suggested by Kalde et al. (2019) who demonstrated binding of TRAPPIII to the vacuole-related RabG3f protein.

Our particular interest in this study has focused on the *Arabidopsis thaliana* TRAPPC8 subunit. In preliminary studies, we have isolated plant lines with mutations in all four TRAPPIII-specific subunits (C8, -11, -12, -13), and of these we have chosen *trappc8* mutants for analysis due to their most pronounced phenotypes. For the yeast and animal counterparts of TRAPPC8, some functional analyzes have been performed. The yeast Trs85p protein has homology to animal and plant TRAPPC8 over the N-terminal half of the protein, but lacks the C-terminal part that interacts with C11 and the C12-C13 dimer. It has long been known that yeast TRAPP functions in ER-to-Golgi transport (Sacher et al., 1998, 2001), and that *trs85*Δ cells also have a defect in the Cvt pathway (*cytoplasm-to-vacuole targeting*, a selective, autophagy-related transport pathway that delivers cytoplasmic cargo directly to the vacuole where it is destined to perform its functions) and impaired (though not completely blocked) autophagy (Meiling-Wesse et al., 2005; Nazarko et al., 2005; Lynch-Day et al., 2010). A function in the secretory pathway at the ER-to-Golgi stage, in Golgi organization, and in autophagy has also been later shown for metazoan TRAPPC8 (Lamb et al., 2016; Scrivens et al., 2011; S. Zhao et al., 2017), C11 (Stanga et al., 2019) and C13 (Ramírez-Peinado et al., 2017). In particular, the C8 subunit has been shown not only to contain the Rab1-binding site (Galindo et al., 2021), but also a membrane-binding site (Harris et al., 2021). Since activation of a specific Rab can take place only on the correct membrane, where its activity is needed, this makes TRAPPC8 a key subunit for the GEF activity of the complex. Additionally, animal C8 and C12 have been implicated in ciliogenesis and the functioning of cilia (Schou et al., 2014; C. Zhang et al., 2020), while TRAPPC11 has been shown to play a role in protein glycosylation (DeRossi et al., 2016; Matalonga et al., 2017; Larson et al., 2018; Munot et al., 2021). Together, these reports indicate a wide variety of functions for TRAPPIII components in the cells of higher eukaryotes.

For plant TRAPPC8, there is less data available. It has been found in post-Golgi, TGN/EE vesicles (Drakakaki et al., 2012; Rosquete et al., 2019b), where it colocalized with core TRAPP components, the TRAPPC11 subunit, and the RabD2 GTPases. Song et al. (2020) have isolated and analyzed *Arabidopsis thaliana* insertion mutant lines with mutations in the AtTRAPPC8-encoding gene (At5g16280). They observed defects in root development and morphology and therefore looked at cellular-level changes in root cells, finding mislocalization of the auxin efflux carrier PIN1 as well as of the TGN/EE proteins RabD2A and VTI12, and abnormalities in the morphology and integrity of compartments of the secretory and endocytic pathways: the trans-Golgi network, early endosomes, and vacuoles.

In this work we have analyzed *Arabidopsis thaliana trappc8* mutants in a broader context. First, by means of the yeast two-hybrid system, we have positioned TRAPPC8 in the network of intracomplex interactions with other TRAPPIII subunits and we found strong analogy to the architecture of the metazoan complex. Next, we have performed in-depth phenotypic characterization of *trappc8* plants, establishing a role for this protein in flower and seed development. Finally, we looked at specific cellular processes known to be affected by *trappc8* mutations in other organisms and we show the engagement of *Arabidopsis* TRAPPC8 in ER functioning and autophagy as well as dolichol turnover.

## METHODS

### Plant lines and growth conditions

In this work *Arabidopsis thaliana* plants of the Col-0 ecotype were used as the WT control line. The analyzed *trappc8* mutants were derived from lines obtained from the NASC collection: SALK_124093 (*trappc8-1*) and SALK_130580 (*trappc8-2*, previously described as *dqc-3* by Song et al., 2020). Primers used for genotyping of the SALK lines are given in Table S1. The same primers were also used to sequence the PCR products resulting from the genotyping procedure, which allowed us to determine precisely the localization of the T-DNA insertions in these lines.

*trappc11* mutants were obtained from the NASC collection. The line referred to as *trappc11-6* was derived from SAIL_118F07 and *trappc11-7* was derived from WiscDSLoxHS012_05B (both previously described as *attrappc11/rog2-*6 and *attrappc11/rog2-7* by Rosquete et al., 2019b). The line *nbr1* was a gift from dr Anna Wawrzyńska (IBB PAS, Warsaw, Poland; described as KO1 in Tarnowski et al., 2020). The line harboring an integrated transgene *pUBI::mCherry-ATG8E* in the WT background was a gift from Dr. Jiwen Liang (Chinese University of Hong Kong, Hong Kong, China; described in Hu et al., 2020), and the line harboring *P_UBI_::mCherry-ATG8E* in the *atg5* background, as well as the lines *atg5* itself (derived from SAIL_129_B07) and *atg2-2*, *atg9-3*, *atg11-1*, and *atg13* were described previously by Stephani et al. (2020). Lines *trappc8-1 [mCherry-ATG8E]* and *trappc8-2 [mCherry-ATG8E]* were obtained by crossing of the lines *trappc8-1* and *-2* (♀) to pollen from the line harboring *[mCherry-ATG8E]* (♂). All primers used for genotyping are listed in Table S1.

For greenhouse culture, plants were grown in soil under standard long day (16 h light/8 h dark) conditions at 22°C. For plate assays, seeds were sown on ½ MS medium supplemented with vitamins, solidified with 1.3% agar, stratified for 2 days and grown vertically in a growth chamber at 22°C, with 70% humidity and a 16 h light/8 h dark period. Where indicated, tunicamycin (Sigma, T7765) was added to the medium from a 1 mg/ml stock in DMSO to a final concentration of 80, 100 or 120 ng/ml. For biochemical experiments, seedlings were grown in liquid ½ MS medium supplemented with 1% sucrose for 2 weeks on a rotary shaker. Conditions used for the starvation sensitivity assays and for confocal microscopy are given below.

### Plasmid construction

The *A. thaliana* TRAPP genes used in the yeast two-hybrid assay (encoding the subunits AtTRAPPC8 (At5g16280), -C11 (At5g65950), -C12 (At4g39820), -C13 (At2g47960), -C2 (At1g80500), -C2L (At2g20930)) were cloned from cDNA prepared either from leaves (-C8, -C13, -C2, -C2L) or flowers (-C11, -C12) of WT plants. The coding sequences were amplified by PCR with appropriate primer pairs (Table S1) and cloned either into the pDONR201 vector by recombination, using the Gateway BP Clonase II Enzyme Mix (Invitrogen), or into the pENTR/d-topo vector (Invitrogen) according to the manufacturer’s instructions. Correct orientation and nucleotide sequence of the products were verified by sequencing. Next, the DNA fragments were transferred by recombination into the appropriate yeast two-hybrid vectors, using the LR Clonase II Kit (Invitrogen). For N-terminal fusions with the GAL4-activation or -binding domains the vectors pGADT7-GW and pGBKT7-GW (gift from Yuhai Cui, Addgene plasmids #61702, 61703; (Q. Lu et al., 2010)) were used, for C-terminal fusions we used pGADCg and pGBKCg (gift from Peter Uetz, Addgene plasmids #20161, 20162; (Stellberger et al., 2010)). All constructs were confirmed by sequencing. The *Escherichia coli* strain used for plasmid propagation was DH5α.

### Yeast two-hybrid (Y2H) assay

The yeast strain used as host for the Y2H assay was AH109 (James et al., 1996). The appropriate plasmids were transformed into yeast cells by the LiAc-PEG method (D. C. Chen et al., 1992), either consecutively or simultaneously. Apart from the test hybrid pairs, each construct was also co-transformed into yeast cells together with the appropriate empty vector (allowing for the expression of the GAL4-AD or –BD alone), to serve as a control of background growth. All transformations were confirmed by colony PCR (primers are listed in Table S1). Cells harboring the correct plasmid pairs were cultured over night (ON) at 28°C in liquid SD (synthetic defined) medium lacking leucine and tryptophan (–leu –trp, supplemented only with adenine and histidine), diluted to a density of 2 OD_600_ units/ml, and used to prepare a series of three 10-fold dilutions. From each dilution series 3 µl-drops were placed on –leu –trp plates (as a control of cell viability and density) and on –leu –trp –his plates (to assay reporter activation), cultured at 28°C for 12 days, and photographed. Each transformed strain was assayed twice.

### Plant morphological observations

For morphological experiments seeds were stratified 2 days and cultured in the greenhouse. Rosette diameter was measured for 4-week old plants as the largest diameter across the rosette. For leaf shape, the largest leaf and the fifth or sixth leaf were collected from 7-week old rosettes. For shoot number, 9-week old plants were used, and each shoot tip (both primary and secondary) was counted. Silique size was judged by taking the three largest of the still green siliques from the main shoot of 9-week old plants. The same siliques were then opened manually with forceps so that their seed content could be visualized.

Flowers were observed in 9-week old plants. Whole inflorescences were removed from the dominant shoot and photographed, then for each inflorescence all open flowers from youngest to oldest were photographed (flower 1 was the first flower from the top that had fully developed, visible petals). Flowers representing stages 14-16 (according to the nomenclature introduced by Smyth et al., 1990) were opened manually with forceps and photographed. Five inflorescences per genotype were observed.

In the case of plate growth assays, the seedlings were cultured on vertical plates, photographed at the indicated time points, and analyzed using the Image J software (v1.53f51). Each experiment was performed twice with similar results.

### Ruthenium red staining

Seeds were imbibed in sterile water for 2 h at RT, stained for 1 h in a 0.02% ruthenium red solution, placed on slides in a drop of water and viewed under the microscope. The photos were analyzed using the Image J software (v1.53f51). The experiment was performed twice with similar results.

### Scanning electron microscopy

For scanning electron microscopy observations, seeds or pollen were spilled directly on microscope tables with double-coated carbon tape, then coated with a thin layer of gold with the use of a sputter coater (Polaron SC7620), and examined using a scanning electron microscope LEO 1430VP (Carl Zeiss).

### In vitro *germination of pollen grains*

Pollen *in vitro* germination was conducted as described in Boavida & McCormick (2007), open flowers from 6-week old plants were used, the germinating grains were viewed under a microscope after 18 h of incubation at 22°C. The experiment was performed twice with similar results.

### Aniline blue staining of pollinated pistils

Arabidopsis Col-0 plants were grown for 5.5 weeks and flowers were manually emasculated. The pistils were hand-pollinated with pollen from the indicated lines, left for 10 h for germination, fixed overnight (ON) in Carnoy’s solution (60% ethanol, 30% chloroform, and 10% acetic acid), and stained with a procedure slightly modified from that of Lu (2011). The fixative was changed to 70% ethanol, then 50% ethanol, 30% ethanol, and water (10 min at RT in each). Then the specimens were moved to 1 M NaOH and left covered ON at RT. The pistils were washed with water for 10 min and stained with 0.1% aniline blue in 50 mM KH_2_PO_4_+K_2_HPO_4_ (pH 7.7) for 3 h in the dark. They were mounted onto microscopic slides in the same phosphate buffer supplied with 50% glycerol and observed under an Eclipse E800 microscope (Nikon Instruments) equipped with a CCD Hamamatsu monochromatic camera. Pollen tube growth (defined as the distance from the pistil stamen to the end of the furthest reaching pollen tube) was measured using the Image J software (v1.53f51) and expressed in relation to the full length of the pistil.

### Starvation sensitivity assays

For carbon starvation assays, the seedlings were grown at standard conditions on solid ½ MS media with 1% sucrose for 7 days and then transferred to plates without sucrose and kept in the dark for another 9 days.

For nitrogen starvation, the seedlings were grown at standard conditions on solid ½ MS media with 0.5% sucrose for 7 days, transferred to plates without nitrogen (½ MS basal salt micronutrient solution (Sigma, M0529) with 3 mM CaCl_2_, 1.5 mM MgSO_4_, 5 mM KCl, 1.25 mM KH_2_PO_4_, 0.5% mannitol and 3 mM MES, pH 5.6) and kept under the same growth conditions for 14 days.

Starvation sensitivity was judged by the extent of leaf chlorosis. The experiment was performed twice with similar results.

### Protein content measurement

Plant material was ground in liquid nitrogen using a mortar and pestle. For assaying 2-week old seedlings grown in liquid medium, exactly 100 mg of ground tissue was supplemented with 300 µl of the following buffer: 50 mM Tris-HCl, pH = 8, 150 mM NaCl, 0.1% Igepal CA630, 10% glycerol, 2.5 mM EDTA, 1 mM PMSF and protease inhibitors (Ultra Easy Pack, Roche), and extraction was allowed to proceed for 2 h at 4°C with shaking. For assaying 4-week old rosette leaves, 30 mg of ground tissue were supplemented with 240 µl of 0.1 N NaOH and extraction was performed for 30 minutes at RT with agitation. In both cases, cell debris was removed by centrifugation for 10 minutes at 4000 g, 4°C, the supernatant was collected and protein content was measured using the Bradford Assay (BioRad), with the appropriate buffer as control. Each measurement was run in three biological replicates.

### Western blotting

Protein extracts for Western blotting were prepared from 100 mg of ground plant material following the same procedure as for protein content measurement. The extracts were mixed with equal amounts of 2x Laemmlie buffer and heated 5 minutes at 95°C. SDS-PAGE was performed on 10% acrylamide gels. 20 µg of protein were loaded per well for anti-NBR1 blots, 15 µg for anti-SKU5 blots, and 25 µg for anti-mCherry. Blotting onto nitrocellulose membranes (Amersham) was conducted using a wet transfer blotting system (BioRad) and controlled by staining with Ponceau S (0.5% in 3% trichloroacetic acid). Membranes were blocked with 3% skimmed milk in TBS + 0.1% Tween-20 (TBST) for 1 hr at RT (or ON at 4°C). The primary antibodies were diluted 1:2000 in TBST with 3% skimmed milk and used for membrane incubation at RT for 1.5-2 h: rabbit polyclonal antibody anti-NBR1 was from Agrisera (AS14 2805), rabbit anti-SKU5 primary antibody was a kind gift from Dr. J.C. Sedbrook (Illinois State University, USA), rabbit polyclonal anti-mCherry from Invitrogen (PA5-34974; used at 1:3000). The goat anti-rabbit-IgG-HRP secondary antibody (Sigma, A0545) was used at a concentration of 1:4000, 1h, RT. TBST was used for membrane washing. Detection was with the SuperSignal West Pico Plus Chemiluminescent Substrate kit (Thermo Scientific) and exposition of Amersham Hyperfilm MP films.

### Confocal microscopy

Wild-type and mutant seedlings carrying the [P_UBI_::mCherry-ATG8E] transgene were cultured in a growth chamber for 5 days on horizontal plates with solid ½ MS medium supplemented with vitamins and 1% sucrose. The seedlings were then transferred to 6-well plates, where 10 seedlings each were incubated 2.5-3 h in the dark at RT with agitation in 3 ml of liquid medium (same as above) supplemented with either 3 µl of 1 mM Concanamycin A (Sigma, C9705) dissolved in DMSO or with 3 µl DMSO alone (control). The seedlings were then mounted on microscopy slides and root cells from the elongation zone were viewed. Imaging of the mCherry fluorophore was carried out on an inverted microscope (Nikon, TE-2000E) with a confocal laser-scanning mode EZ-C1. mCherry fluorescence was excited with light emitted by a Green He-Ne Laser set at 30% (1.0 mW, 543 nm; Melles Griot, USA) then collected with a 605/75 nm single band pass emission filter (BrightLine, Semrock) and displayed in false red. Cells have been imaged using a 60x oil immersion objective (Nikon, CFI Plan Apochromat NA 1.4). All confocal adjustments (the pinhole diameter set to 30 µm, the gain of the detector set to 8.4, the time dwell set to 10.08 µs) were the same during the experiment. Confocal sections were collected at 0.5 µm intervals, resulting in stacks encompassing the fixed volume of: 120 µm x 75 µm x 20 µm. Puncta of diameter 0.4 to 2.5 µm were counted manually from the obtained image stacks. For each genotype/condition 8-16 image stacks taken from 4-9 individual seedlings were scored. The full experiment was repeated twice with similar results.

### Real-time quantitative PCR (RT-qPCR)

100 mg of plant material ground in liquid nitrogen was used for RNA isolation with the GeneJET Plant RNA Purification Mini Kit (Thermo Scientific). The samples were digested with the RapidOut DNA Removal Kit (Thermo Scientific), and 0.5 μg of RNA was reverse-transcribed using the RevertAid First Strand cDNA Synthesis Kit (Thermo Scientific). The obtained cDNA was quantified by qPCR using SG qPCR Master Mix (2×) plus ROX Solution (EURx, Gdansk, Poland) and a StepOnePlus Real-Time PCR System (Applied Biosystems). The cDNA was diluted 10-fold and 10 μl was used in a total reaction volume of 25 μl per well. All primers used for RT-qPCR analyses are listed in Table S1. The gene encoding PROTEIN PHOSPHATASE 2A SUBUNIT A3 (PP2A) was used as internal reference. The expression of each gene was examined in two technical and three biological replicates. The relative expression levels were determined using the 2^−ΔΔCt^ method and normalized to expression in WT plants.

### Polyisoprenoid isolation from Arabidopsis tissue

Plant material (collected from several plants; ca. 1 g of fresh mass, except the experiment with lyophilized seedlings where 200–400 mg of dry material were used) was either ground in liquid nitrogen and transferred to 20 ml of a 1:1 (v/v) chloroform/methanol mixture (C/M), or homogenized directly in 20 ml of the same mixture with an Ultra-Turrax apparatus (IKA Labortechnik). The samples were supplemented with 10 µg of Prenol-14 as internal standard and agitated for 48 h at room temperature. The C/M extracts were filtered and evaporated under a stream of nitrogen. The obtained lipids were resuspended in 5 ml of hydrolysis mixture (7.5% (w/v) KOH in ethanol : toluene : water, 4:5:1 by volume) and incubated for 1 h at 96°C. After cooling, 5 ml of each water and hexane were added and the mixture was vortexed. After phase separation, the organic phase was collected, and extraction of the aqueous phase was repeated twice with 5 ml hexane each. The pooled organic phases were evaporated under a stream of nitrogen and dissolved in 500 µl of hexane. The extracts were loaded onto 2 ml-silica gel columns in hexane and eluted with 10 ml 2% diethyl ether in hexane, and 20 ml of 15% diethyl ether in hexane. The latter fraction was evaporated and dissolved in 100 µl of 2-propanol; all of the sample was used for HPLC/UV analysis. Three independent biological replicates were run for each experiment.

### HPLC/UV analysis of polyisoprenoid alcohols

Polyisoprenoid alcohols were analyzed by HPLC/UV as described previously (Jozwiak et al., 2013), with modifications. Runs were performed on a 4.6×75 mm ZORBAX XDB-C18 (3.5 μm) reversed-phase column (Agilent) using a Waters dual-pump apparatus, a Waters gradient programmer, and a Waters Photodiode Array Detector (spectrum range: 210–400 nm). The chain length and identity of lipids were confirmed by comparison with external standards of a polyprenol mixture (Prenol-9–25) and dolichol mixture (Dolichol-17–21). Quantitative determination of polyisoprenoids was performed using the internal standards. Integration of the HPLC/UV chromatograms was performed with the Empower software (Waters). All polyprenol and dolichol standards were from the Collection of Polyprenols, Institute of Biochemistry and Biophysics, Polish Academy of Sciences (Warsaw, Poland).

### Statistical analysis

Statistical analysis was performed with GraphPad Prism 9.5.1 (GraphPad, San Diego, CA, USA) software. For inheritance analysis, the χ2 test or the binomial test were performed against Mendelian H_0_ hypotheses, as described in Table 2. For analysis of root length, seed axis length or pollen tube length, the structures were measured using ImageJ and compared pair-wise using unpaired t-tests with Welch’s correction against the H_0_ hypotheses that the lengths are equal. Protein and isoprenoid content as well as RT-qPCR results were compared pair-wise using unpaired t-tests against the H_0_ hypotheses that the content is unchanged. The numbers of mCherry puncta in the autophagy experiment were analyzed with unpaired t-tests with the Welch’s correction. The box-and-whiskers plots used in this paper show min to max values with median indicated and with whiskers representing the 25^th^ and 75^th^ percentile, the bar graphs display mean values with standard deviations.

## RESULTS

### Binary interactions in the plant TRAPPIII complex are consistent with the structure of metazoan TRAPPIII

The structure of the plant TRAPPIII complex has not been investigated, but the subunit composition has been established (Kalde et al., 2019; Rosquete et al., 2019b) and resembles that of metazoan TRAPPIII, whose structure was published recently (Galindo et al., 2021). To see if the complexes can be considered structurally similar, we investigated binary interactions among the TRAPPIII-specific (C8, C11, C12, C13) and adaptor (C2, C2L) subunits from *Arabidopsis thaliana* using the yeast two-hybrid system.

We concentrated on the interactions of the two large subunits, TRAPPC8 and C11. For these proteins, we prepared both N-terminal and C-terminal fusions with the GAL4-activation domain (AD) and GAL4-binding domain (BD) and tested all possible binary interactions with N-terminal AD and BD fusions of the C2, C2L, C12 and C13 subunits. Consistently with the metazoan structure (Galindo et al., 2021), which shows that the N-terminal parts of the C8 and C11 subunits are involved in interactions with the adaptor proteins, the C-terminal fusions displayed clear interactions with the C2 and C2L subunits (Fig. 1A): C8-C2 and C11-C2L, while the N-terminal fusions could not interact with the adaptor subunits. When assayed together, the C8 and C11 subunits did not interact with each other in any combination (not shown); it is possible that their interaction requires the presence of the subunits C12 and/or C13.

**Fig. 1.**
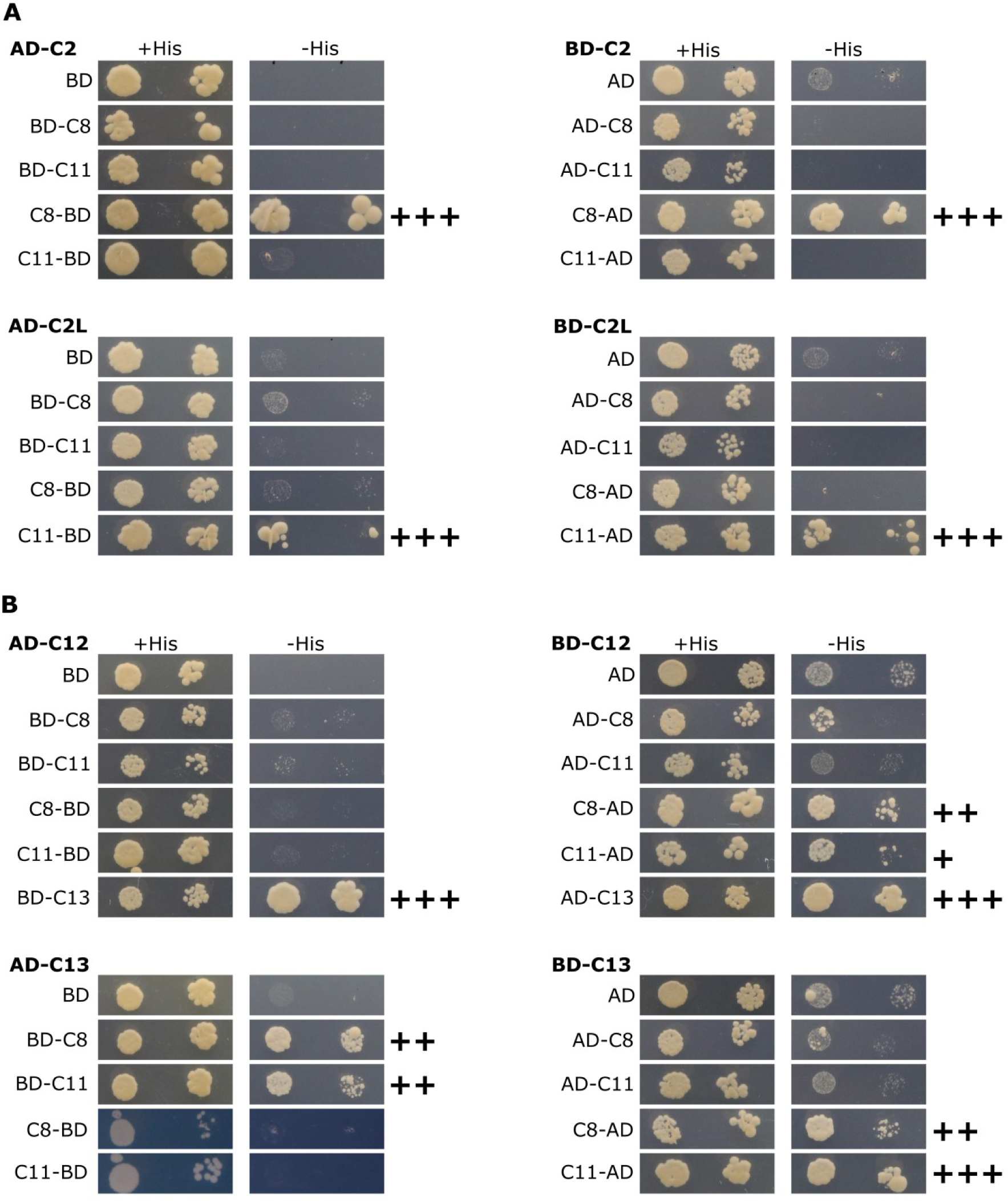
Binary interactions between selected TRAPP subunits assayed using the yeast two-hybrid system. For each panel, one hybrid protein is indicated above the panel and the other to the left. Abbreviations used: AD – Gal4 activation domain, BD – Gal4 DNA-binding domain. The names of TRAPP subunits are shortened to subunit numbers. +His marks control plates, - His are test plates showing reporter activation. Interactions scored as positive (when compared to the relevant control interactions) are indicated by plus signs next to the images. (A) Interactions of TRAPPC8 and TRAPPC11 with the adaptor subunits TRAPPC2 and TRAPPC2L. (B) Interactions of TRAPPC12 and TRAPPC13 with each other and with the large subunits TRAPPC8 and TRAPPC11.

The C12 and C13 subunits interacted strongly with each other, in all combinations, confirming the presence of a C12-C13 dimer. Their interactions with the large subunits presented a less clear picture: as BD-fusions, both subunits interacted with the C-terminal C8 and C11 fusions, but as AD fusions only the C13 subunit showed an interaction with N-terminal C8 and C11 (Fig.1B). This suggests that C13 is likely mediating this interaction and that a free C-terminus is not crucial for the interaction of the C8 and C11 subunits with the C12-C13 dimer. Taken together, these results indicate that the general structure of plant TRAPPIII is similar to that of the metazoan complex.

### Loss of Arabidopsis TRAPPC8 leads to smaller plants with excessive branching and defective siliques

To learn more about the *in vivo* functions of plant TRAPPIII we decided to investigate *A. thaliana* mutants defective in TRAPPC8, the only subunit that is both specific for this complex and has a (partial) homolog in yeast. We surveyed two T-DNA insertion lines with mutations mapped in *TRAPPC8*: SALK_124093 and SALK_130580.

Segregation of the mutant alleles did not follow the 1:2:1 pattern of inheritance (Table 1). For both mutants, an analysis of 48 plants derived from parental heterozygous lines showed that significantly less homozygotes and less heterozygotes were obtained, despite the fact that the parental lines showed no morphological abnormalities. We then crossed manually the heterozygous lines with wild-type plants, to see which parent displayed the defects (Table 1). The results were similar for both alleles: when pollen was derived from wild-type plants, segregation followed the Mendelian inheritance pattern, but when pollen was derived from heterozygous plants, disproportionately few mutant plants were obtained, defining a clear male transmission defect for *trappc8* mutations.

**Table 1.** Segregation of the *trappc8-1* and *trappc8-2* alleles.

| Cross | Expected ratio for Mendelian segregation | Observed ratio | Number of observations | *P*-value | Significance |
| --- | --- | --- | --- | --- | --- |
| **self-pollination** | WT:het:hom | WT:het:hom |  |  |  |
| *trappc8-1*<sup>+/-</sup> | 1 : 2 : 1 | 15 : 31 : 2 | 48 | 0.0038* | ** |
| *trappc8-2*<sup>+/-</sup> | 1 : 2 : 1 | 21 : 25 : 2 | 48 | 0.0005* | *** |
| **manual crossing** | WT:het | WT:het |  |  |  |
| *trappc8-1*<sup>+/-</sup> ♀ × WT ♂ | 1 : 1 | 71 : 58 | 129 | 0.2541** | NS |
| WT ♀ × *trappc8-1*<sup>+/-</sup> ♂ | 1 : 1 | 118 : 10 | 128 | <0.0001** | **** |
| *trappc8-2*<sup>+/-</sup> ♀ × WT ♂ | 1 : 1 | 67 : 65 | 132 | 0.8624** | NS |
| WT ♀ × *trappc8-2*<sup>+/-</sup> ♂ | 1 : 1 | 90 : 9 | 99 | <0.0001** | **** |

\* Analyzed with the $\chi^2$ test against the $H_0$ hypothesis that segregation is Mendelian.
\*\* Analyzed with the binomial two-tailed test against the $H_0$ hypothesis that segregation is Mendelian.

Still, in both cases we were able to isolate homozygous lines; the location of the insertion sites was confirmed by PCR and sequencing (Fig. S1A, B). Both mutants produce truncated transcripts of the gene, but lack the full-length version (Fig. S1C, D). We termed them *trappc8-1* (SALK_124093, carrying an insertion in intron 20) and *trappc8-2* (SALK_130580, insertion in intron 10; this is the same mutant as the *dqc-3* mutant described by Song et al., 2020).

Next we characterized the overall morphology of both mutant plant lines. Data published previously on the *trappc8-2/dqc-3* line included only observations of root phenotypes in seedlings (Song et al., 2020), here we looked also at entire plants. Both *trappc8* lines grew slower and were smaller than wild-type plants, they had smaller rosettes with pointed, more serrated leaves, kinked stem growth, and an excessive secondary branching phenotype (Fig. 2A-C, E; for stem growth see Fig. 5A). Bolting time remained unchanged, but the flowering stage and silique development were prolonged. The plants produced small siliques (Fig. 2F) which were partially or completely empty (Fig. 2G). The mutants, though not completely sterile, produced very low amounts of seeds per plant (Fig. 2D). When cultured on plates, mutant seedlings showed slower root growth as well as premature leaf bleaching (Fig. 3).

**Fig. 2.**
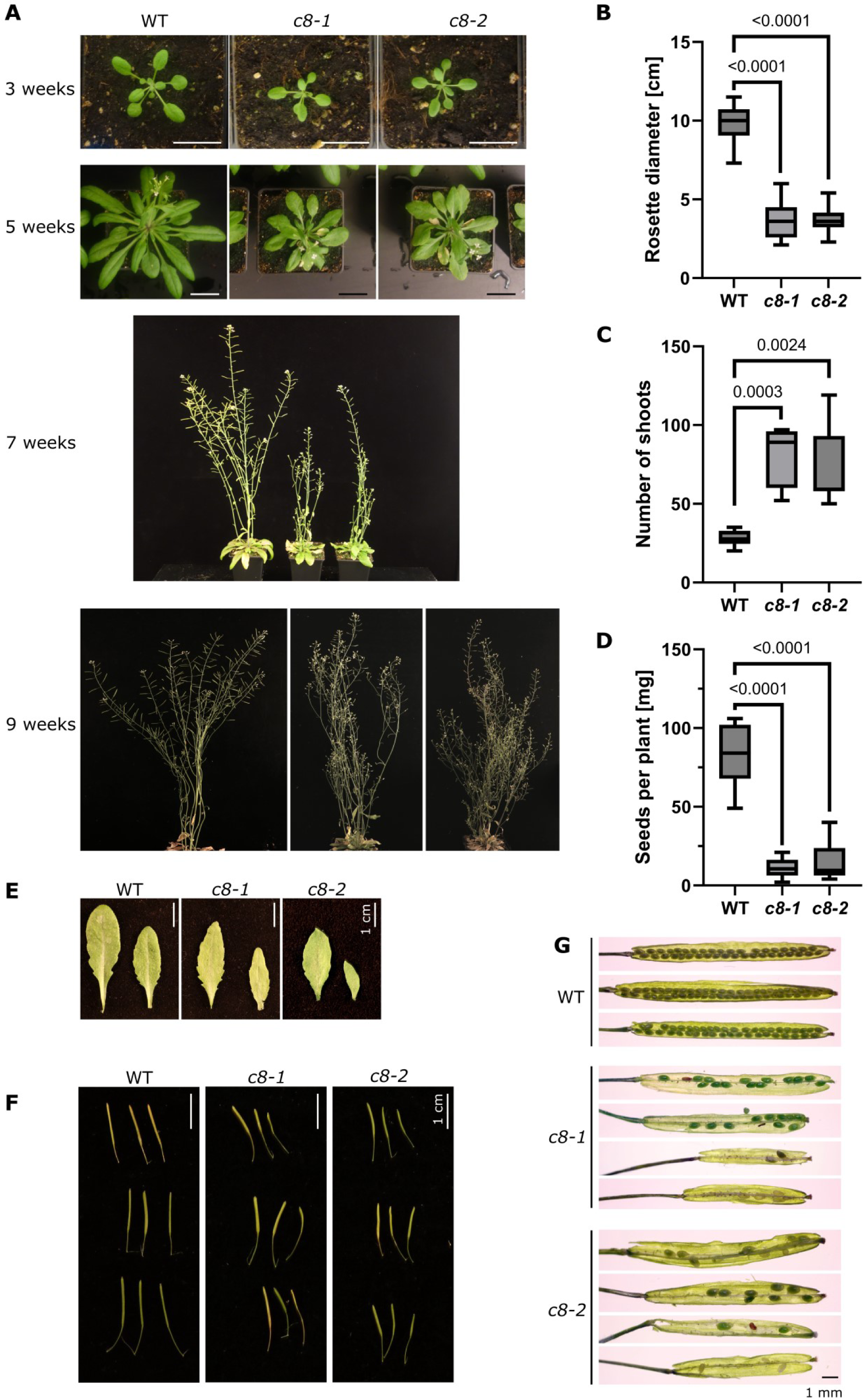
Phenotypic analysis of the mutant lines *trappc8-1* and *trappc8-2*. (A) Overall morphology of WT and mutant plants cultured in soil. Bar: 3 cm (B) Quantification of rosette size of 5-week old plants. 20 plants were measured for each genotype. The P values of unpaired t-tests with Welch’s correction are indicated on the graph (also in C and D). (C) Quantification of total number of shoots (primary and secondary) of 9-week old plants. 6-7 plants were scored for each genotype. (D) Quantification of amount of seeds produced per plant. The seeds of 8-14 individual plants were weighed for each genotype. (E) Representative rosette leaves from 7.5-week old plants of the indicated genotypes. (F) For 3 representative plants each, 3 mature but unopened siliques were photographed. (G) The siliques shown in panel F were opened and photographed, representative examples are shown.

**Fig. 3.**
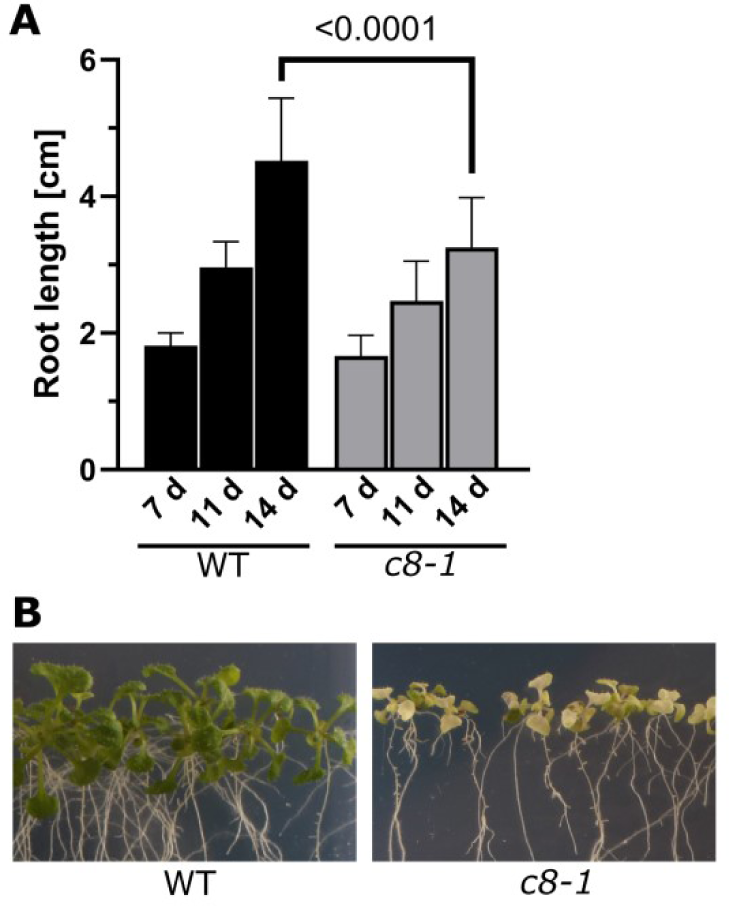
Phenotypes of *trappc8-1* mutant seedlings cultured on agar plates. (A) Seedlings were cultured on vertical agar plates (1/2 MS) and photographed at 7, 11 and 14 days after sowing. Root length was measured using ImageJ. 39-40 seedlings were measured for each time point. The P value of an unpaired t-test with Welch’s correction is indicated on the graph. (B) After 21 days of growth, mutant seedlings show premature leaf bleaching.

**Fig. 4.**
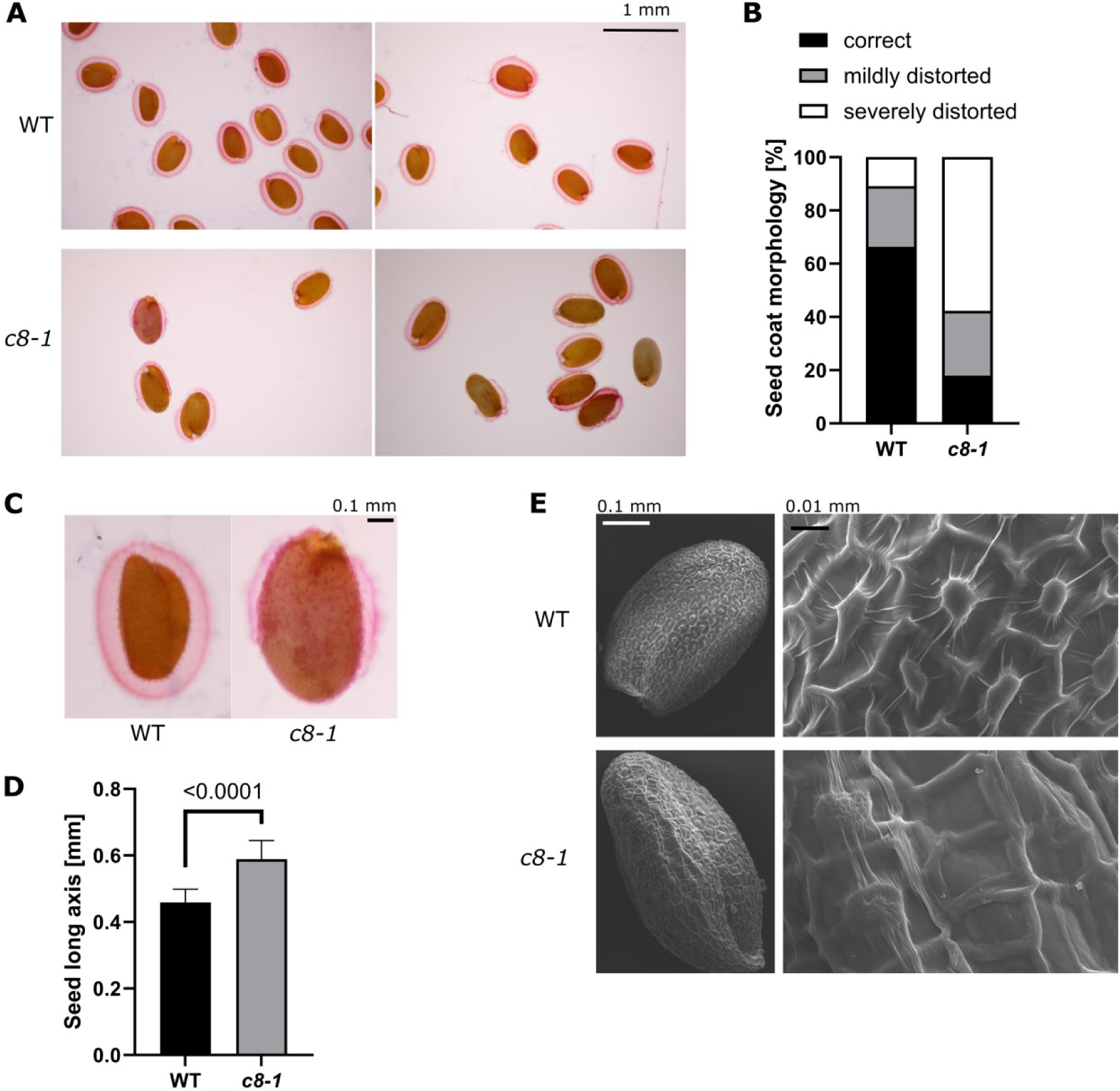
*trappc8-1* mutant plants produce abnormal seeds. (A) WT and *trappc8-1* seeds were imbibed, stained with ruthenium red and viewed under the microscope. (B) Quantification of mucilage coat morphology. Approx. 80 seeds per line were manually assigned to 3 coat morphology categories: correct – the coat fully covers the seed, mildly distorted – the coat thins out / loses continuity in one spot, severely distorted – more parts of the seed are coatless. (C) Close-up view of seeds from A. Mutant seeds are larger and misshaped. (D) Quantification of seed size. The same seeds as scored in B were measured (length of long axis) using ImageJ. The P value of an unpaired t-test with Welch’s correction is indicated on the graph. (E) SEM images of WT and *trappc8-1* seeds. Aberrant morphology of the seed coat epidermal cells is visible.

**Fig. 5.**
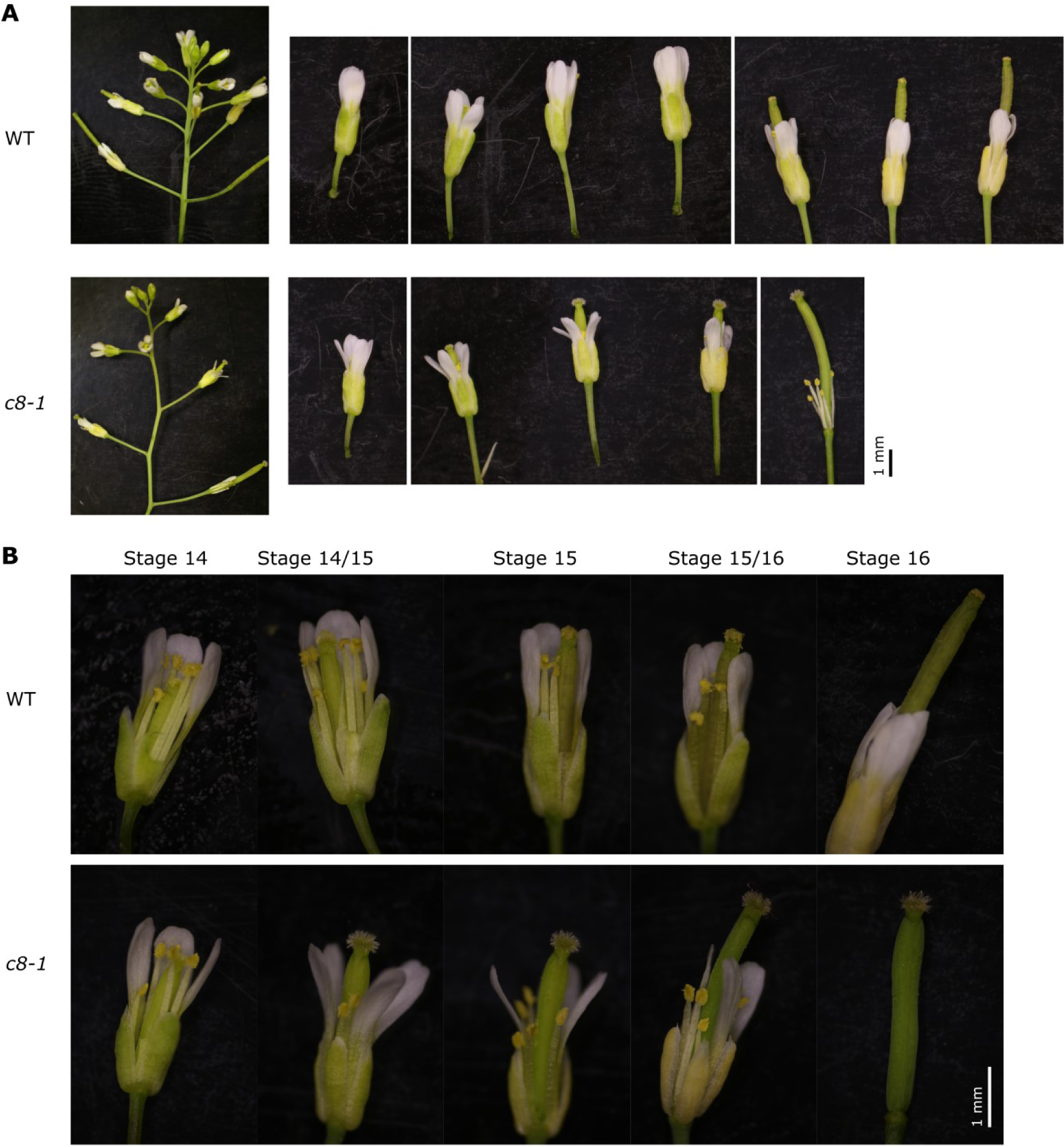
*trappc8-1* mutant plants have defective flower development. (A) A representative inflorescence is shown for WT and *trappc8-1* plants. Note the kinked stem growth of the mutant. To the right, all flowers of the particular inflorescences are shown, starting from the first flower with visible petals. The protruding pistil of the mutant flowers is visible. (B) Close-up of flowers from panel A. Flower development stages as defined by Smyth et al. (1990) are indicated.

These morphological phenotypes remain in contrast to the appearance of the published *trappc11* mutants (Rosquete et al., 2019b) which resembled wild-type plants. We also directly compared the growth of *trappc8* and *trappc11* plants (Fig. S2) and confirmed the major morphological differences. These results suggest that the TRAPPC8 subunit may perform functions additional to those performed together with TRAPPC11, as part of the TRAPPIII complex.

### TRAPPC8 is necessary for the development of seed coat cells and for mucilage deposition

We next inspected microscopically the seeds obtained from the homozygotic *trappc8* mutants. We observed imbibed seeds using ruthenium red staining which visualizes acid polysaccharides. This method allowed us to assess not only seed size and shape, but also the morphology of the mucilage layers deposited on the seed outer surface. Seed morphology was visibly changed. The mutant seeds were larger and more elongated and had thinner mucilage layers (Fig. 4A-D). The mucilage layers were often severely distorted, with large parts being very thin or completely absent. We visualized the seed outer surface by scanning electron microscopy (SEM) and we could see that the morphology of seed coat epidermal cells was defective (Fig. 4E). The epidermal cells were irregular and had distorted columellae, suggesting defects in deposition of the secondary cell wall. Despite the observed defects, the seeds were viable: when sowed on plates, they germinated only slightly worse than wild-type seeds (92% compared to 99%). The observed seed phenotypes are similar to those noted previously for exocyst mutants (Kulich et al., 2010) and are consistent with the involvement of TRAPPIII in secretory processes.

### The trappc8 mutant has impaired flower development and pollen functioning

The defective siliques prompted us to inspect the flowers of the *trappc8* mutant. We noticed that although initial stages of flower development proceed normally (up to stage 13 (Smyth et al., 1990)), later the pistil outpaces the stamens and petals (Fig. 5). This leads to a situation where the mature stigma of stage 14 – which is functional in the mutants, as shown by the successful manual pollination (Table 1) – remains without contact with the dehiscing anthers and receives very little pollen, or none at all, and eventually dries out. This effect could account for the very low seed yield of the homozygous mutants, but the observed allele frequencies in the progeny of *trappc8* heterozygous lines (Table 1) led us to investigate also pollen functioning.

Indeed, when we observed pollen grains by SEM, we found that they displayed visibly altered morphology (Fig. 6). Overall morphology of the grains was impaired, with about half of the grains misshaped. The structure of the pollen grain surface was also changed: the pollen cell wall structure was disturbed, in some regions of the grains even completely unformed. During pollen development, the pollen cell wall components are synthesized and secreted by tapetum cells (Ma et al., 2021), so the process is dependent on correct functioning of the secretory pathway in these cells.

**Fig. 6.**
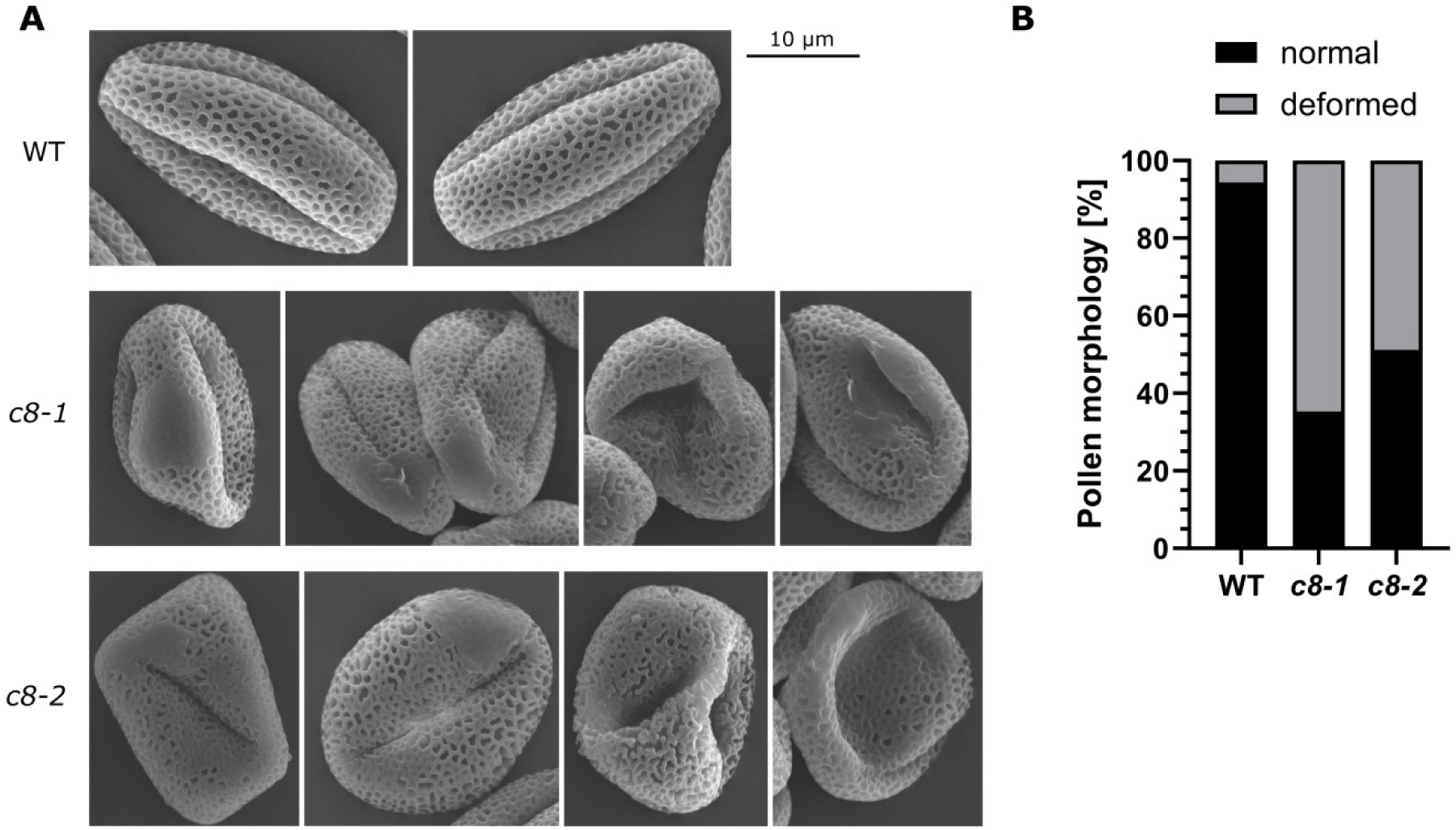
*trappc8* mutant flowers produce aberrant pollen grains. (A) SEM images of WT and *trappc8* pollen grains. (B) Quantification of photographed pollen. 130-170 grains were scored per genotype.

We also reasoned that secretory pathway disturbances could lead to problems with pollen tube growth, which is highly dependent on secretion. However, when *trappc8* pollen grains were germinated *in vitro*, they were capable of growing tubes morphologically similar to those of wild-type pollen (Fig. 7A). We next performed hand-pollination of emasculated wild-type flowers to observe the germination of *trappc8* pollen grains on wild-type pistils. Here, we could clearly see that mutant pollen was able to adhere to the stigma and to germinate, and that the pollen tubes successfully made their way through the style (Fig. 7B), but did not reach as far down the transmitting tract as wild-type pollen tubes after 10 hours (Fig. 7C), showing that they grow slower. Incidents where we could see a mutant pollen tube that left the transmitting tract and directed its growth towards an ovule were seldom when compared to wild-type pollen tubes (Fig. 7D). In some cases, the growth of the mutant pollen tubes inside the pistil seemed chaotic (Fig. 7E). These observations showed that the defect in secretion in the *trappc8* mutants is not so strong that it would prevent pollen tube growth (although it may be slowing it down), but rather that the ability of *trappc8* pollen to respond to signals emitted by the female gametophytes might be impaired. In particular, defects seem to occur at the stage of funicular guidance (Lopes et al., 2019), since the pollen tubes do not efficiently turn from the transmitting tracts towards the ovules.

**Fig. 7.**
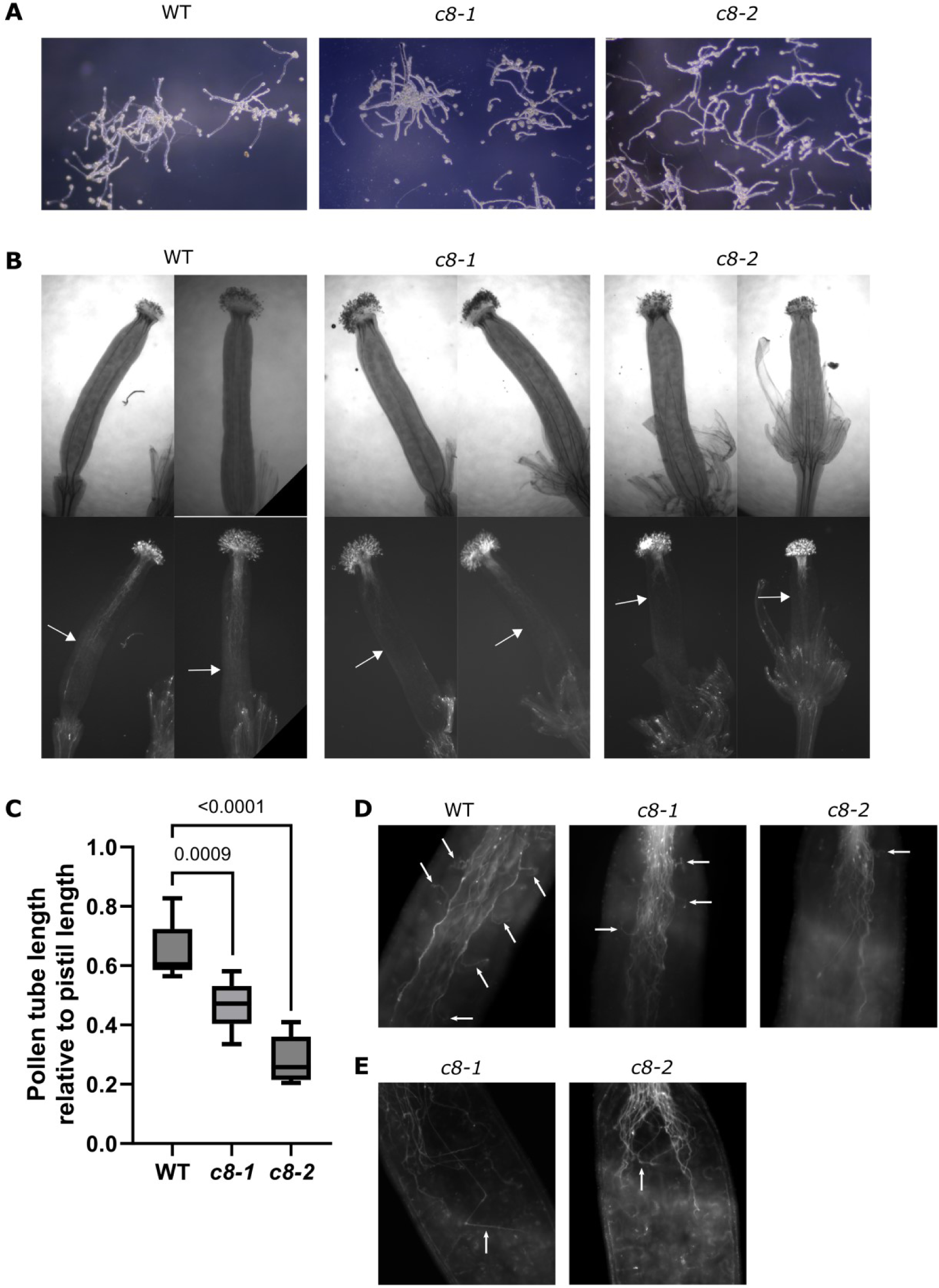
*trappc8* pollen tubes are functionally deficient. (A) WT and mutant pollen grains were germinated *in vitro* and photographed. (B) WT and mutant pollen grains were placed manually on emasculated WT pistils, allowed to germinate for 10 h, fixed, stained with aniline blue and viewed under the microscope. Arrows mark the furthest reaching pollen tube for each pistil shown. (C) Quantification of experiment shown in B. For each line, 8-9 pistils were scored. The P values of unpaired t-tests with Welch’s correction are indicated on the graph. (D) Close-up views of representative stained pistils. Arrows mark instances where a pollen tube can be seen turning towards an ovary. (E) Examples of mutant pollen tubes growing chaotically inside the pistils.

The low fertility of the *trappc8* lines probably arises as a sum of the observed defects: the aberrant flower morphology (discrepancy between pistil and stamen size), disturbed pollen biogenesis, and impaired ovule-targeting competence of the male gametophyte.

### Arabidopsis TRAPPC8 is involved in autophagy

The yeast homolog of TRAPPC8, Trs85p, is an established factor in autophagy (Meiling-Wesse et al., 2005; Nazarko et al., 2005), and we hypothesized that Arabidopsis TRAPPC8 might also play a role in this process. To test this, we first assayed the starvation sensitivity of the mutants, since defects in autophagy typically lead to worse survival under starvation conditions. For all autophagy-related experiments, we used an *atg5* mutant line as positive control: the ATG5 protein is involved in lipidation of ATG8 and is indispensable for autophagosome formation. We compared the growth of *trappc8* mutant seedlings with that of wild-type and autophagy-deficient *atg5* seedlings under conditions of nitrogen and carbon starvation. Judged by the extent of leaf chlorosis occurring, the *trappc8* mutants were slightly sensitive to nitrogen starvation, though not to carbon starvation (Fig. 8A). However, the effect was very mild and should be viewed in the context of the phenotype of premature leaf chlorosis displayed by the *trappc8* mutants (see Fig. 3), so we looked for further data. Another general feature of autophagy-deficient plant mutants is a mild increase in total protein content per fresh weight (Guiboileau et al., 2013), resulting from defects in the turnover of protein aggregates. The *trappc8* mutants did show a slight increase in protein content, but in our hands even the control line *atg5* showed such a minor increase that this phenotype was difficult to assess (Fig. S3). We therefore further tested autophagic degradation also biochemically, by assaying the level of NBR1, a cargo receptor for selective autophagy in plant cells (Zhang & Chen, 2020). In *trappc8* seedlings, as in *atg5* seedlings, NBR1 protein content was strongly elevated in comparison to wild-type levels, while the corresponding mRNA was induced only approximately 2-fold – likely not enough to account on its own for the strong protein accumulation. This was consistent with the notion that autophagic degradation does not proceed efficiently in the mutants (Fig. 8B,C).

**Fig. 8.**
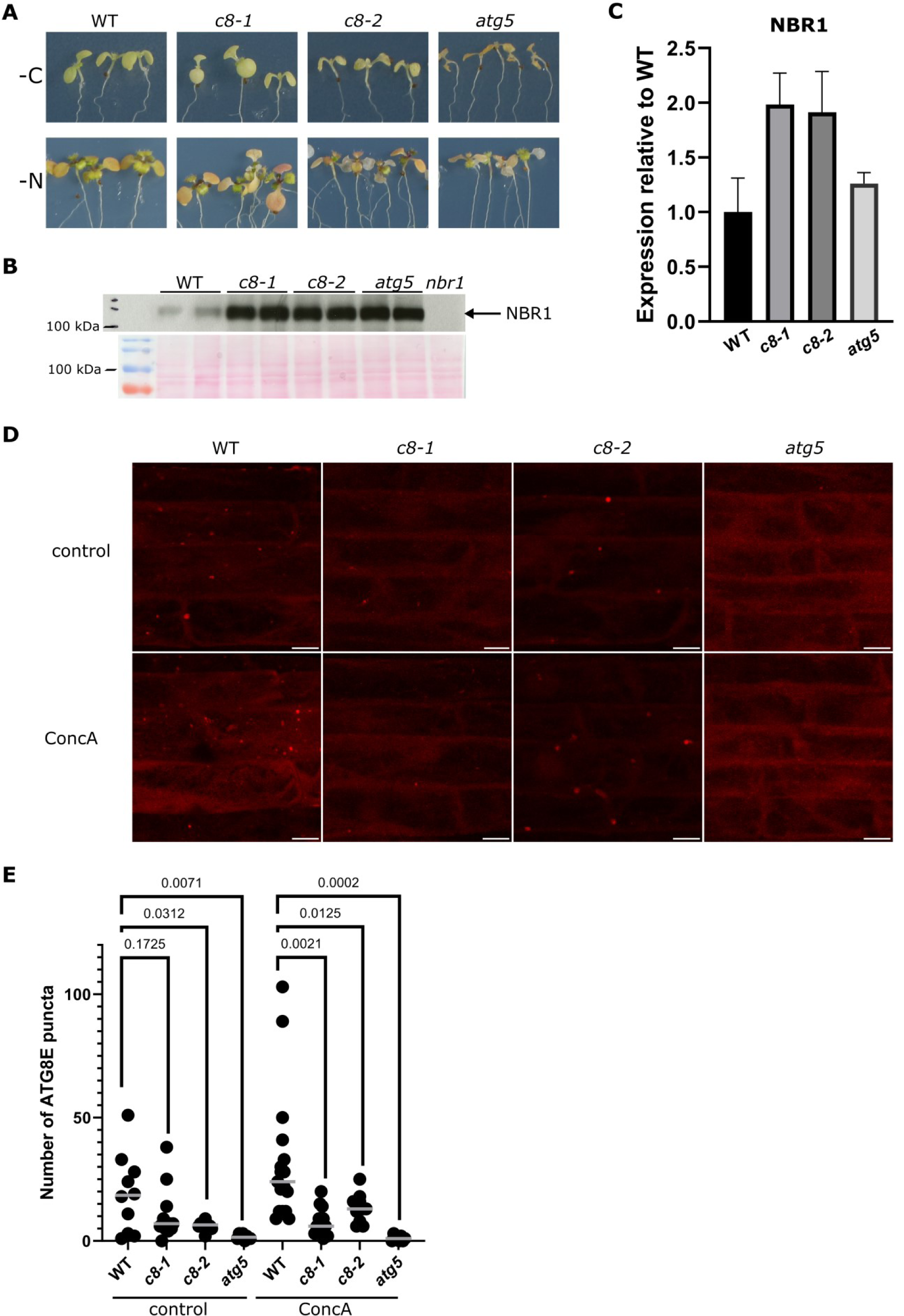
*trappc8* mutants display mild autophagy defects. (A) 7-day old seedlings of the indicated genotypes were transferred to conditions of either carbon (-C) or nitrogen (-N) starvation and observed for leaf chlorosis. (B) Plate-grown 2-week old seedlings of the indicated genotypes were subjected to protein extraction and Western blot analysis with an anti-NBR1 antibody. Ponceau S staining of the membrane is shown below to demonstrate gel loading. (C) Liquid-grown 2-week old seedlings of the indicated genotypes were subjected to RNA extraction and RT-qPCR analysis. The levels of transcripts for the *NBR1* gene are shown in relation to the level for WT seedlings. (D) 5-day old seedlings expressing the fusion protein mCherry-ATG8E in the indicated genetic backgrounds were incubated for 2.5-3 h with or without ConcA and root epidermal cells were photographed in a confocal microscope with fixed settings. Representative images are shown. The image stacks were flattened for presentation. Scale bar: 10 µm. (E) mCherry-stained puncta were counted manually in the image stacks. The horizontal bars indicate the median for each result set. The numerical values on the graph are P values for unpaired t-tests with Welch’s correction.

Finally, to monitor autophagic flux directly, we constructed *trappc8-1* and *-2* lines expressing a fluorescent version of ATG8E (P_UBI_::mCherry-ATG8E). The ATG8E protein remains associated with autophagosomes from the stage of pre-autophagosomal membrane expansion until full autophagosome maturation and vacuole delivery, and it is an established autophagosome marker in Arabidopsis cells (Contento et al., 2005). Expression of the fusion protein was confirmed by Western blotting with an anti-mCherry antibody (Fig. S4). Since the mutants showed only minor starvation sensitivity, but a large increase in NBR1 levels under conditions that were not autophagy-inducing, we decided to assay basal autophagy. The notion that basal autophagy would be impaired in our mutants was also consistent with previous results from yeast cells, where TRAPPIII was found necessary for autophagy under nutrient-rich conditions (Shirahama-Noda et al., 2013). 5-day old seedlings expressing the *mCherry-ATG8E* transgene in WT, *trappc8* or *atg5* backgrounds were incubated with or without Concanamycin A (ConcA) ─ an inhibitor of the vacuolar H^+^-pump which prevents proper acidification of the vacuole and thus the degradation of autophagosomes in its lumen ─ and then observed by confocal microscopy to assess the number of autophagosomes per given region in root epiderm (Fig. 8D,E). Both *trappc8* mutants showed decreased numbers of autophagosomes compared to WT seedlings, confirming that autophagy is not proceeding properly in these cells. At the same time, in the presence of ConcA there was a difference in phenotype penetration observable between the *atg5* and the *trappc8* mutants: in *atg5* cells there were almost no ATG8E-positive dots at all, while in *trappc8* cells some puncta were observed. This suggests that TRAPPC8 might play a regulatory role in autophagy rather than being structurally indispensable for the process.

### Induction of the UPR in trappc8 mutants

Defects in protein trafficking through the early secretory pathway may cause protein accumulation in the ER and lead to ER stress (Pastor-Cantizano et al., 2018). If TRAPPIII would be required for trafficking at earlier steps than TGN/EE compartments, such as ER exit and ER-to-Golgi trafficking, then *trappc8* mutations could cause ER stress and lead to induction of the unfolded protein response (UPR). To investigate this, we assayed the *trappc8* mutants for transcriptional upregulation of several UPR markers. These included the ER chaperones: CNX1 (calnexin 1, recognizes misfolded transmembrane and glycosylated proteins (Bloemeke et al., 2022, and references therein)), BiP2 and BiP3 (chaperones for soluble ER lumen proteins) and their J-domain interactor ERDJ3A (Pobre et al., 2019), and the protein disulfide isomerase PDI6 (also known to participate in recognizing misfolded clients in the ER lumen (Lu & Christopher, 2008)). We included also analysis of the bZIP60s transcript – an unconventionally spliced version of the mRNA for the transcription factor bZIP60 – which is induced by UPR and IRE1 activation (Nagashima et al., 2011). All of these are established markers of the UPR in *A. thaliana* (Pastor-Cantizano et al., 2018). As shown in Fig. 9, transcriptional induction was very strong for BiP3 and significant for ERDJ3A; for PDI6 a mild increase was observed. Also the bZIP60s mRNA was significantly induced. Together, these results show that the *trappc8* mutants have constitutive activation of the UPR.

**Fig. 9.**
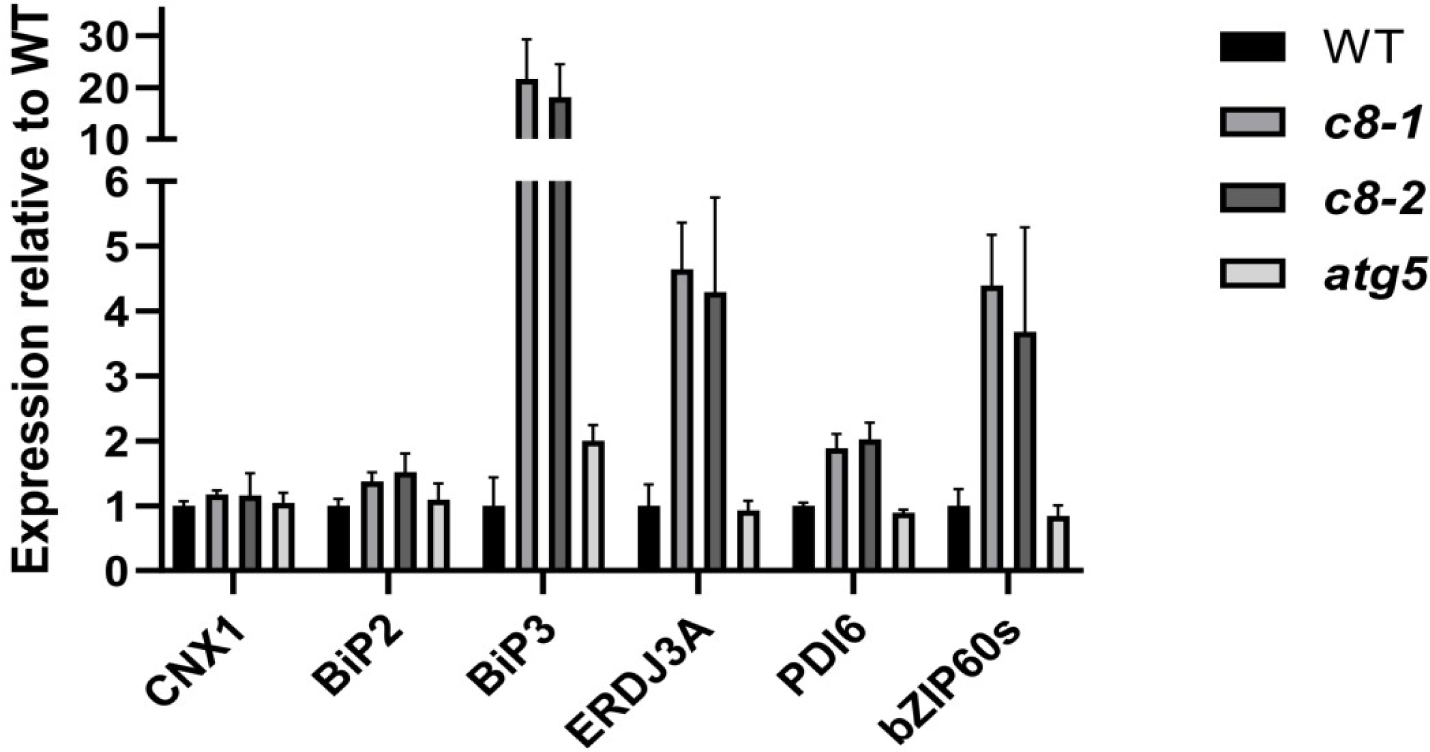
*trappc8* mutants, but not the *atg5* mutant, display a constitutive ER stress response. RNA extracted from seedlings of the indicated genotypes grown for 2 weeks in liquid medium was used for RT-qPCR with primers detecting six UPR marker transcripts.

To rule out the possibility that ER stress in the analyzed mutants results from the observed defects in autophagy, which could impair the cell’s ability to clear protein aggregates accumulating in the ER, we included the *atg5* mutant in the analysis (Fig. 9). The mutant did not display any significant UPR induction, which was consistent with previous data for the *A. thaliana atg5* mutant (in a transcriptomics analysis no induction was found for UPR genes; Havé et al., 2019) as well as with data from yeast cells (mutations impairing autophagy did not lead to an induction of the UPR unless they were accompanied by an overproduction of ER proteins; Lipatova et al., 2020). This suggests that the constitutive ER stress observed in *trappc8* is not a secondary effect of autophagy deficiency, but results rather from direct defects in ER functioning.

### Effects of TRAPPC8 loss on protein glycosylation and dolichol levels

The established function of the TRAPPIII complex as a GEF for Rab1/Ypt1 in animal and yeast cells (Galindo et al., 2021; Harris et al., 2021; Joiner et al., 2021) suggests a role of this complex in early secretory transport and autophagy. In plants, TRAPPIII has been implicated also in some additional vesicular transport routes in the cell: *trappc8* mutants showed defects in endocytosis and vacuolar transport (Song et al., 2020), while results obtained for *trappc11* mutants showed a role of TRAPPIII in the late secretory pathway, at a post-Golgi stage (Rosquete et al., 2019b). On the other hand, in zebrafish embryos and HeLa cells, mutation of the *TRAPPC11* subunit gene has been shown to cause defects in protein glycosylation and lowered levels of lipid-linked oligosaccharide (LLO) precursors (DeRossi et al., 2016). In that work the effect was not visible after silencing of the gene for the TRAPPC8 subunit, but the expression of the targeted protein was only partially suppressed under the applied experimental conditions. We reasoned that some of the phenotypic traits described here correlated well with a potential defect in protein glycosylation: defects in communication between the pollen tube and female gametophyte have been previously described for *N*-glycosylation-deficient mutants (Lindner et al., 2015) and for GPI-anchoring-deficient mutants (Capron et al., 2008), and the flower and pollen grain morphology of *trappc8* plants resembled that of *pprd2* mutants, which are deficient in the biosynthesis of dolichol, an indispensable cofactor of glycosylation (Jozwiak et al., 2015). We therefore decided to look at protein glycosylation and dolichol levels in *trappc8* plants.

First we tested extracts from wild-type, *trappc8-1* and *-2* seedlings with antibodies against the specific protein SKU5, which is known to be both strongly glycosylated and decorated with a GPI anchor and can thus serve as a marker of glycosylation defects in *A. thaliana* (Sedbrook et al., 2002), but neither a change in gel migration nor lowered levels of SKU5 were detected (Fig. S5).

In animal cells, the *trappc11^-^* mutation was synthetically lethal with treatment with tunicamycin (Tun) (DeRossi et al., 2016) – an inhibitor of *N*-glycosylation. We investigated the sensitivity of *trappc8* seedlings to Tun treatment using a plate growth assay. The mutants showed a mild root growth defect when cultured on regular media (Fig. 3A, Fig. 10), but when Tun was added, the WT seedlings were affected much more visibly than the mutant seedlings, and at the highest Tun concentration used (120 ng/ml) the mutant seedlings had significantly longer roots than WT seedlings (Fig. 10A). Thus, to our surprise, loss of full-length TRAPPC8 actually conferred Tun resistance.

**Fig. 10.**
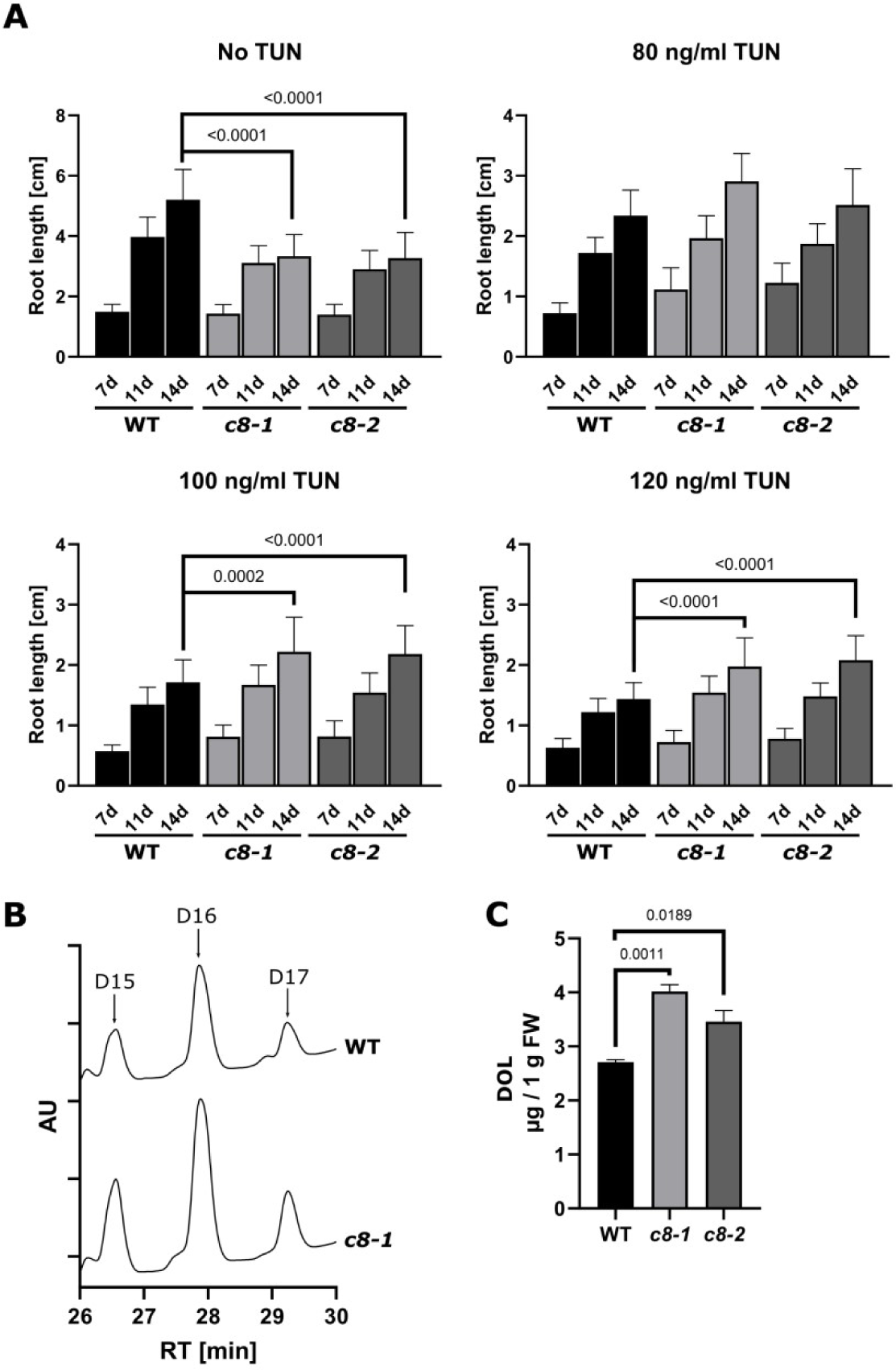
*trappc8* mutant plants show tunicamycin resistance and increased dolichol accumulation. (A) WT and *trappc8* seeds were sowed on plates containing the indicated concentrations of tunicamycin (TUN) in the growth medium. Seedlings were photographed after 7, 11 and 14 days of growth and root length was measured using ImageJ. The values given on the graphs are P values for unpaired t-tests with Welch’s correction. (B) WT and mutant seedlings were grown for 2 weeks in liquid medium, collected and subjected to isoprenoid isolation. The obtained samples were analyzed by HPLC-UV. Representative chromatograms are shown, with peaks corresponding to dolichol-15 (D15), dolichol-16 (D16) and dolichol-17 (D17) marked. AU – arbitrary units, RT – retention time. (C) Quantification of dolichol accumulation. The sum of dolichol-15, -16 and -17 is plotted. FW – fresh weight. The numerical values on the graph are P values for unpaired t-tests with Welch’s correction.

From these results we concluded that in the absence of TRAPPC8, protein glycosylation is conducted efficiently but the glycosylation process is changed in some aspects, leading to Tun insensitivity. To gain more insight into this phenomenon, we measured directly the intracellular levels of free dolichol (Dol) in *trappc8* seedlings cultured in liquid medium. Both mutant lines showed an increase in dolichol content up to 130-150% of wild-type values (Fig. 10B). The results were similar in lyophilized seedlings, showing that they were not resulting from differences in water content (Fig. S6A). The Dol content was elevated also in rosette leaves of 4-week old plants cultured in soil, confirming that this was a mild but reproducible effect, not connected to culture conditions (Fig. S6B). This result was surprising, but it was in line with the Tun resistance, since dolichol phosphate (Dol-P) is one of the substrates of the enzyme GlcNAc-1-P transferase, which is the direct target of Tun.

To find out if the additional dolichol results from an induction of its biosynthesis, we checked if the transcription of enzymes responsible for dolichol biosynthesis was upregulated in the mutants (Fig. S7). The transcription of the MVA and MEP pathway genes: *HMGR1*, *HMGR2*, *MVK*, *DXS*, *DXR*, *MCT*, as well as of two FPP synthase genes: *FPPS1* and *FPPS2*, and of genes for the two final enzymes of Dol biosynthesis: *CPT3* (Gawarecka et al., 2022) and *PPRD2* (Jozwiak et al., 2015), remained at wild-type levels. Slight, but non-significant, induction was observed only for *EVN* which encodes a dolichol kinase (Lindner et al., 2015). These results suggest that dolichol biosynthesis is not increased in the *trappc8* context.

We considered then the possibility that elevated Dol levels result from reduced rates of dolichol degradation, but no specific mechanism of Dol degradation in cells has been described yet. We reasoned that, since dolichol resides in the ER membrane and the ER is likely an important source of membrane for autophagy (Zhen & Stenmark, 2023), a decrease in autophagic membrane turnover could be the underlying cause of decreased Dol degradation. To test this idea, we measured Dol levels in several homozygous *atg* mutants of *A. thaliana*: *atg2*, *atg5*, *atg9*, *atg11*, *atg13*. However, these mutants, though autophagy-deficient, mostly showed no significant increase in Dol levels (Fig. S8A); only *atg2* showed a mild accumulation (up to 130%) similar to the effect visible in *trappc8* lines. We also tested the effects of Tun on the growth of a representative *atg* mutant line (*atg5*) and we did not observe any Tun resistance effect (Fig. S8B). These results show that the Dol increase observed in our mutants is specific for *trappc8*, not a general effect of autophagic deficiency.

## DISCUSSION

In this study we build on the known structure of the metazoan TRAPPIII complex to propose an initial map of plant TRAPPIII. The binary interaction data we provide shows that the general architecture of the plant complex is similar to that of metazoan TRAPPIII: TRAPPC8 and C11 interact with the adaptor subunits C2 and C2L, respectively. Further, we confirm that TRAPPC13 is part of the complex (as indicated before by the results of Kalde et al., 2019) and that it remains in a strong interaction with TRAPPC12, unlike the yeast protein Trs65p (closest homolog of TRAPPC13) which is part of TRAPPII (Choi et al., 2011).

We then set out to learn more about the functions of the plant TRAPPIII complex by characterizing one of its key subunits, TRAPPC8. We performed a careful phenotypic analysis of entire mutant plants. Our results are in line with the possible functions of TRAPPC8 (and likely the whole TRAPPIII complex) in sorting and recycling at the TGN/EE and in vacuolar traffic, but they also add new potential sites of action for the protein.

### TRAPPC8 and Rab regulation

The developmental defects observed in this study can largely be linked to misfunctions in Rab regulation and, as a result, in protein sorting and secretion. Previous work has shown that aberrant protein sorting in *trappc8* root cells causes the auxin efflux carrier PIN1 to be partly mislocalized to the vacuole, leading to an aberrant auxin response which was associated with defects in root growth and gravitropism (Song et al., 2020). Here we observe multiple further developmental defects, apart from retarded root growth: small size of plants, excessive branching, serrated leaves, kinked stem growth, delayed flowering, and pronounced fertility defects resulting in small siliques. These phenotypes can be explained by disturbances in hormone homeostasis and in delivery to the plasma membrane of integral or secreted components, pointing to a role of TRAPPC8 in the secretory pathway.

Auxin mutants display similar flowering and fertility phenotypes (Tan et al., 2023), though in this case the seeds produced are smaller than WT, while *trappc8* mutants produce larger seeds. Incorrect sorting and secretion of proteins may also underlie the changes in seed coat epidermal cells and in seed mucilage deposition (Kulich et al., 2010) as well as the abnormal morphology of *trappc8* pollen grains, because efficient secretion of components by tapetum cells is necessary for formation of the exine layer (Zhao et al., 2016; Aboulela et al., 2018). The defects in pollen tube guidance may be related both to impaired delivery of membrane-bound receptors (Guan et al., 2014; Stührwohldt et al., 2015) and to impaired ER functioning (Wang et al., 2008; Lu et al., 2011).

Finally, the *trappc8* phenotypes strongly resemble those of *rabD2* mutants. The double mutant *rabD2b rabD2c* was described to have short siliques, defects in pollen grain morphology (with cell wall defects similar to those of *trappc8* pollen), decreased pollen tube growth rate, and a reduced seed set (Peng et al., 2011). The pollen grain cell wall defects were also similar in *mon1* mutants (Cui et al., 2017), which are defective in the activation of RabG proteins. RabG proteins localize to vacuolar and prevacuolar membranes where they regulate traffic coming both from the endocytic and autophagic routes (Kwon et al., 2013; Cui et al., 2017). Also Rab prenylation mutants, which have partial defects at multiple inner trafficking routes, display similar abnormal flower morphology and pollen phenotypes (Gutkowska et al., 2015; Gutkowska et al., 2021). Therefore, the phenotypic data for *trappc8* mutants supports the involvement of the protein in Rab regulation at more than one site. Interestingly, *trappc11* mutants do not display the same morphological and fertility phenotypes as the *trappc8* lines – a phenomenon requiring further investigation.

### TRAPPC8 and ER functioning

Some of our results point to a requirement for TRAPPC8 in ER homeostasis and functioning. First, the analyzed *trappc8* mutants have constitutive activation of the UPR response, suggesting an accumulation of misfolded and/or accumulated proteins in the ER. We confirm that this defect is not a secondary effect of autophagy impairment, but an autophagy-independent phenomenon. Second, we describe for the first time a possible defect in the degradation or removal of dolichol, an ER lipid serving as a cofactor for protein glycosylation. Overall levels of dolichol are elevated in *trappc8* mutants while no transcriptional induction in the dolichol biosynthesis pathways can be observed. We reasoned that this might result from decreased dolichol turnover, e.g. through a decreased rate of lipid flow into autophagosome membranes. Interestingly, of the *atg* mutants tested, only the line deficient in the lipid transfer protein ATG2 showed a somewhat similar dolichol phenotype. The increase in dolichol levels is (consistently) accompanied by tunicamycin resistance of *trappc8* seedlings, and accordingly, protein glycosylation proceeds efficiently. This observation is in line with a recent report describing an Arabidopsis Golgi trafficking mutant (*cog7*) which displayed enhanced N-glycosylation in the Golgi apparatus (elevated transcript levels of glycosyltransferase genes) (Choi et al., 2023), but it remains in contrast to observations from animal cells, where mutations in a different TRAPPIII subunit, TRAPPC11, led to decreased levels of LLO (lipid-linked oligosaccharides) and glycosylation defects (however, dolichol levels were not tested in any animal models) (DeRossi et al., 2016; Matalonga et al., 2017; Larson et al., 2018; Munot et al., 2021). We directly tested dolichol levels in two *A. thaliana trappc11* mutants (Fig. S9) and confirmed that in these lines no decrease in dolichol levels occurs (though no consistent increase either). The mechanism linking TRAPPC11 with changes in the glycosylation process in animal cells awaits explanation. Similarly, the role of TRAPPC8 in maintaining correct regulation of dolichol levels in plant cells needs further investigation. Taken together, these data suggest a link between TRAPPC8 and ER homeostasis in eukaryotic cells.

### TRAPPC8 and autophagy

We also tested the efficiency of autophagy in our mutants and we found that the process is impaired. Under conditions where autophagy was not induced, we saw a significant decrease in the number of autophagosomes being formed by cells. On the other hand, overall autophagic flux in induced (starved) cells was not dramatically low, as shown by the weakness of the starvation survival phenotypes of the mutants.

That it would be basal autophagy that would be impaired in our mutants is consistent with previous results obtained from yeast cells, where TRAPPIII was found necessary for Atg9p cycling from early endosomes to the ER under nutrient-rich conditions, while the requirement was by-passed upon starvation (Shirahama-Noda et al., 2013); also in mammalian cells TRAPPIII subunits were shown to be necessary for ATG9A recycling and for autophagosome formation (Imai et al., 2016; Lamb et al., 2016) or for autophagic flux under selected conditions (Ramírez-Peinado et al., 2017; Stanga et al., 2019). It was shown also in Arabidopsis that the role of ATG9 for autophagosome formation is indispensible (Zhuang et al., 2017) and our data suggest that here, as in yeast, the requirement for TRAPPC8 is largely by-passed under starvation conditions. It is worth noting that in plants ER stress is a known inducer of autophagy (Liu et al., 2012) and our mutants show constitutive ER stress, however, autophagosome formation is not induced in response to this situation, showing that this induction pathway – unlike starvation – does require TRAPPC8 function. Our results support therefore the concept that there are various routes in plant cells for autophagy signalling.

A further interesting question is whether the elevated levels of dolichols observed in *trappc8* cells might directly underlie or otherwise be linked to their autophagy impairment. Lipids have been suggested to act as modulators of autophagy with different outcomes depending on the lipid subclass, the autophagic process involved, and the cell type and age (Fan et al., 2019; Hernández-Cáceres et al., 2024), and the specific mechanisms of this modulation remain elusive.

### TRAPPC8 and senescence

The premature senescence phenotype of *trappc8* mutant plants can be connected both to autophagy dysfunction and dolichol accumulation. *atg* mutants display early leaf senescence in response to various nutrient-limiting conditions (reviewed by Chen et al., 2019). Increasing dolichol levels have long been considered a marker of aging in animal tissues (Chojnacki & Dallner, 1988; Parentini et al., 2005), and in plants dolichol levels are also known to increase with age (Gawarecka et al., 2022). Interestingly, the recent work of Choi et al. (2023) shows that a defect in the functioning of retrograde transport from the Golgi apparatus can be the primary cause of both premature leaf senescence and enhancement of the glycosylation machinery – in line with our results which show similar effects of a primary dysfunction of the ER.

### Conclusions

Results described in this report document that the architecture of the TRAPPIII complex is conserved in plants. Moreover, TRAPPC8 is implicated in the functioning of numerous events of intracellular trafficking, ranging from ER functioning to secretion and autophagy (Fig. 11). This wide spectrum of cellular processes involving TRAPPC8 must result in a complex map of interactors – to be discovered yet. Interestingly, misfunction of TRAPPC8 leads to increased accumulation of dolichol, which in turn protects the cells against deficiencies in protein glycosylation.

**Fig. 11.**
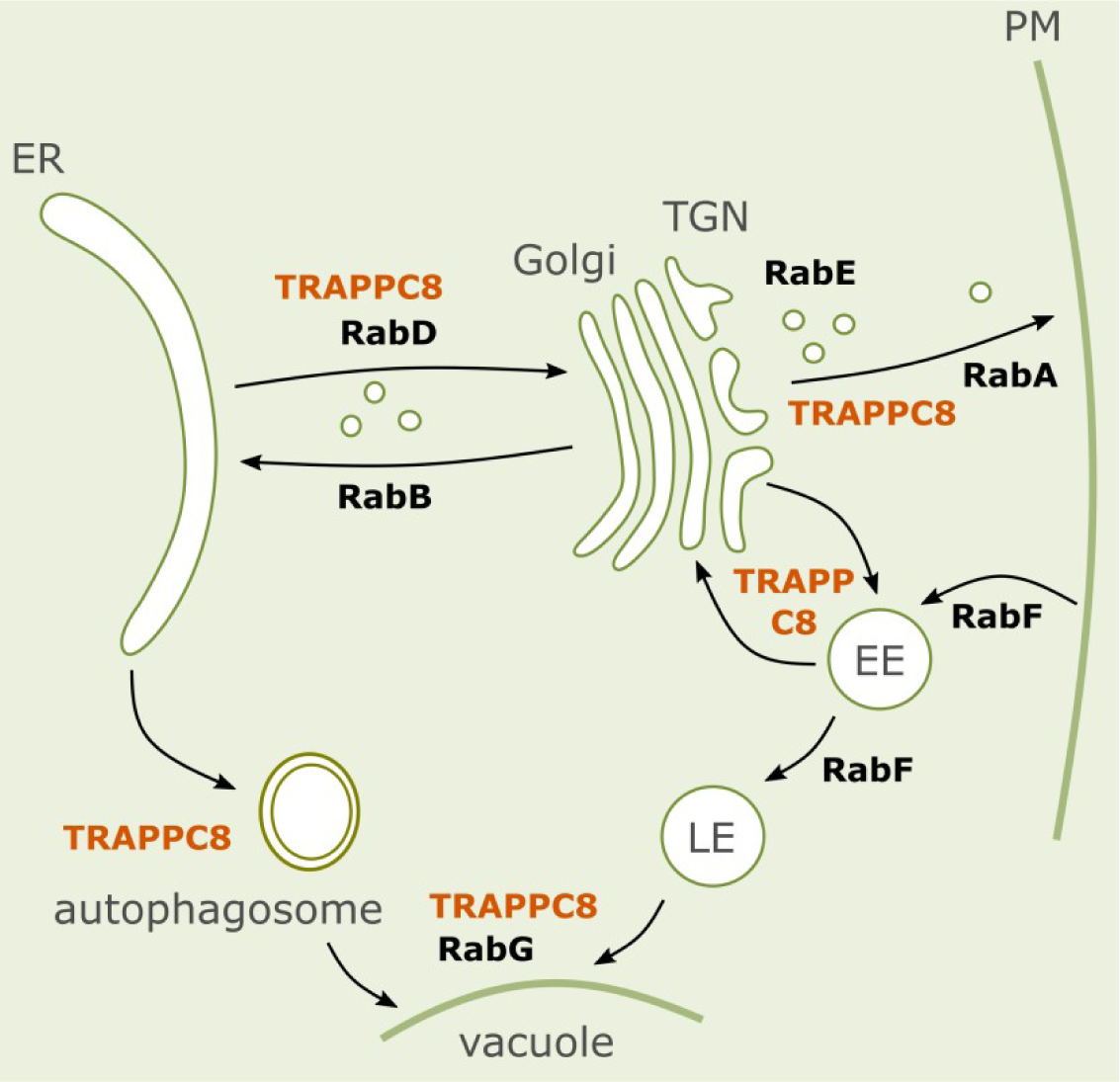
Cellular sites of action of TRAPPC8, a subunit of the TRAPPIII complex. EE – early endosomes, LE – late endosomes. AtTRAPPC8 is involved in regulation of transport to the Golgi and vacuole, as well as in sorting and recycling at the TGN and EE and in the biogenesis of autophagosomes.

## ACKNOWLEDGMENTS

We thank dr Małgorzata Gutkowska for numerous suggestions and comments during the course of this work as well as for critical reading of the manuscript. We thank drs Anna Wawrzyńska (IBB, Polish Academy of Sciences) and Jiwen Liang (Chinese University of Hong Kong) for sharing published plant lines, and dr Daniel Buszewicz (previously: IBB, Polish Academy of Sciences) for sharing RT-qPCR primers and unpublished results of primer tests.

## AUTHOR CONTRIBUTIONS

MHS generated many concepts of the work, performed the experiments and wrote the first version of the manuscript. NP performed part of the RT-qPCR experiments. AAM was responsible for confocal microscopy. JN conducted SEM imaging. YD took part in designing and supervised autophagy-related experiments. ES formulated the project and supervised the work. All authors took part in preparing the final version of the manuscript.

## FUNDING

This work was supported by grants from the National Science Centre of Poland: UMO-2018/29/B/NZ3/01033 to Ewa Swiezewska and UMO-2018/02/X/NZ3/00822 to Marta Hoffman-Sommer.

## Notes

### Competing Interest Statement

The authors have declared no competing interest.

