## Supplemental Table S1 for "The *Arabidopsis thaliana* TRAPPIII subunit AtTRAPPC8/AtTRS85 is involved in ER functioning and autophagy"

**Supplementary Table S1** for

"**The *Arabidopsis thaliana* TRAPPIII subunit AtTRAPPC8/AtTRS85 is involved in ER functioning and autophagy**"

by Hoffman-Sommer et al. (2024)

**Table S1**. Primers used in this study.

| Primer name | Sequence (5’→3’) | Purpose |
|---|---|---|
| S124093L | TGATCAATTGAACCCTCAAGC | Genotyping of SALK_124093 |
| S124093R | TTTCAGAGTCGGTGAAATTGG | Genotyping of SALK_124093 |
| S130580L | ACTTGGATGTGCTCTCACTGG | Genotyping of SALK_130580 |
| S130580R | ATGGTTCACCGCATCACTTAC | Genotyping of SALK_130580 |
| W01205BL | ACCACCACCCACATCAACTAG | Genotyping of WiscDSLoxHS012_05B |
| W01205BR | TTTTCACTTGAAATGGTTGGC | Genotyping of WiscDSLoxHS012_05B |
| I118f07L | GCAACCAATTCGAATTCTTTG | Genotyping of SAIL_118F07 |
| I118f07R | CTGGAATTGAAGAAGAGCACG | Genotyping of SAIL_118F07 |
| atg5-LP | ATTTGCTATTTGTTTGGCACG | Genotyping of SAIL_129_B07 |
| atg5-RP | TACCGTTCATGACAGAGGTCC | Genotyping of SAIL_129_B07 |
| SAIL-LB1 | TAGCATCTGAATTTCATAACCAATCT CGATACAC | Genotyping of SAIL lines |
| LBb1.3 | ATTTTGCCGATTTCGGAAC | Genotyping of SALK lines |
| WDLH-L4 | TGATCCATGTAGATTTCCCGGACATG AAG | Genotyping of WiscDSLoxHS lines |
| mCH-8E | ACGAGTTCGAGATCGAGG | Genotyping of lines with transgene *[mCherry-ATG8E]* |
| ATG8E-R | ATTGAAGAAGCACCGAATGT | Genotyping of lines with transgene *[mCherry-ATG8E]* |
| C8-BP-F | GGGGACAAGTTTGTACAAAAAAGCA GGCTTC**ATG**GTGGAGCCGGTGAACTC TT | Cloning of AtTRAPPC8 into the pDONR201 vector |
| C8-BP-R | GGGGACCACTTTGTACAAGAAAGCTG GGTC**CTC**AGACTGAAGAACAGTAAG G | Cloning of AtTRAPPC8 into the pDONR201 vector |
| C2-BP-F | GGGGACAAGTTTGTACAAAAAAGCA GGCTTC**ATG**ATCGTTTGCGTCGCCGT C | Cloning of AtTRAPPC2L into the pDONR201 vector |
| C2-BP-R | GGGGACCACTTTGTACAAGAAAGCTG | Cloning of AtTRAPPC2L into the |

| | | |
|---|---|---|
| | GGTC**GTT**CAAACCGTACGATCCAAC | pDONR201 vector |
| BP-C2w-F | GGGGACAAGTTTGTACAAAAAAGCA GGCTTC**ATG**GCTAACACTGCCTGC | Cloning of AtTRAPPC2 into the pDONR201 vector |
| BP-C2w-R | GGGGACCACTTTGTACAAGAAAGCTG GGTCCAGGTACTTTCTTGCAAG | Cloning of AtTRAPPC2 into the pDONR201 vector |
| C11pop-F | CACC**ATG**GAGGAATACCC | Cloning of AtTRAPPC11 into the pENTR/d-TOPO vector |
| TR-C11-R | CTTGCTGGTAGAGATGGC | Cloning of AtTRAPPC11 into the pENTR/d-TOPO vector |
| C12pop-F | CACC**ATG**GTATCTATTGGAAAGAC | Cloning of AtTRAPPC12 into the pENTR/d-TOPO vector |
| TR-C12-R | GACTCTGGTACAAGATGAATCAAAG | Cloning of AtTRAPPC12 into the pENTR/d-TOPO vector |
| C13-BP-F | GGGGACAAGTTTGTACAAAAAAGCA GGCTTC**ATG**AGCGCGACGCAGACG | Cloning of AtTRAPPC13 into the pDONR201 vector |
| C13-BP-R | GGGGACCACTTTGTACAAGAAAGCTG GGTC**GTC**TGTCTCTACAAATATCTC | Cloning of AtTRAPPC13 into the pDONR201 vector |
| C8-F3 | CCTCCCAGCCAAACTGATGT | Colony PCR of yeast for Y2H assay and semi-quantitative PCR |
| C8-R4 | GGACTTACCTGAACCGAGGG | Colony PCR of yeast for Y2H assay and semi-quantitative PCR |
| C11-F3 | ACAAGCTCAGCAACTGGTCA | Colony PCR of yeast for Y2H assay |
| C11-R4 | ACAACAACAGTTCCAGGGCA | Colony PCR of yeast for Y2H assay |
| C2L-F3 | GAGGCTTTTCTTGGCTTGCTC | Colony PCR of yeast for Y2H assay |
| C2L-R4 | CTTCCTGAAGAAACTTCTCACATCG | Colony PCR of yeast for Y2H assay |
| C2w-F1 | ACCTAGCATGGACTACAAGTGC | Colony PCR of yeast for Y2H assay |
| C2w-R2 | CATCAGTCGGGTATGGCCTG | Colony PCR of yeast for Y2H assay |
| C12-F3 | GGGACGCATTACAGGTACGA | Colony PCR of yeast for Y2H assay |
| C12-R4 | ACGATCTAACCCTTCCTGACG | Colony PCR of yeast for Y2H assay |
| C13-F3 | TCCAATTGAATCTTATTGCCTCCA | Colony PCR of yeast for Y2H assay |
| C13-R4 | ACTCGGGGAGGCTGTCTTAT | Colony PCR of yeast for Y2H assay |
| C8-F1 | CAACCAAACGTAGAGGTTGCT | RT-qPCR analysis and semi- |

| | | |
|---|---|---|
| | | quantitative PCR |
| C8-R2 | TACCGAAACCATGAGGGTGC | RT-qPCR analysis and semi-quantitative PCR |
| C8-F13 | TGGTTCCATCGTAGGTGTGC | RT-qPCR analysis |
| C8-R14 | CAAGCCTGGGTAGACTCTTGA | RT-qPCR analysis |
| CNX1-F | ATGAGACAACGGCAACTATTTTCC (Cho and Kanehara, 2017) | RT-qPCR analysis |
| CNX1-R | CCATAATCCTCATGTCCTTCACT (Cho and Kanehara, 2017) | RT-qPCR analysis |
| BiP1/2-F | GGTATCGAGACTGTAGGAGG (Cho and Kanehara, 2017) | RT-qPCR analysis |
| BiP1/2-R | GGTACGTTGTGAAAACCTGA (Cho and Kanehara, 2017) | RT-qPCR analysis |
| BiP3-F | CGAAACGTCTGATTGGAAGAA (Cho and Kanehara, 2017) | RT-qPCR analysis |
| BiP3-R | GGCTTCCCATCTTTGTTCAC (Cho and Kanehara, 2017) | RT-qPCR analysis |
| ERDJ3A-F | TCAAGTGGTGGTGGTTTCAACT (Pastor-Cantizano et al., 2018) | RT-qPCR analysis |
| ERDJ3A-R | CCCACCGCCCATATTTTG (Pastor-Cantizano et al., 2018) | RT-qPCR analysis |
| PDI6-F | CGAAGTGGCTTTGTCATTCCA (Pastor-Cantizano et al., 2018) | RT-qPCR analysis |
| PDI6-R | GCGGTTGCGTCCAATTTT (Pastor-Cantizano et al., 2018) | RT-qPCR analysis |
| bZIP60s-F | GGAGACGATGATGCTGTGGCT (Pastor-Cantizano et al., 2018) | RT-qPCR analysis |
| bZIP60s-R | CAGGGAACCCAACAGCAGACT (Pastor-Cantizano et al., 2018) | RT-qPCR analysis |
| NBR1_5 | TGGTCGATACATTTCTTATTGGAGG (Jung et al., 2020) | RT-qPCR analysis |

| | | |
|---|---|---|
| NBR1_6 | GGGAGGCATTAAGGTTCAGTC (Jung et al., 2020) | RT-qPCR analysis |
| HMGR1-F | TCTATCGAGGTGGGGACAGT | RT-qPCR analysis |
| HMGR1-R | GATCGTCGCTAGCCTCCTTG | RT-qPCR analysis |
| HMGR2-F2 | CCCAGCTCAAAACGTGGAGA | RT-qPCR analysis |
| HMGR2-R2 | TCCCACCTCCAACAGTACCA | RT-qPCR analysis |
| MVK-F | ATGTGGCTCTCCTCTGGGAT | RT-qPCR analysis |
| MVK-R | CCAGACCCGTATGGAAGCTC | RT-qPCR analysis |
| DXS-F | GCGTGCTTATGACCAGGTTG | RT-qPCR analysis |
| DXS-R | ATGTGTCGGACCATCAGCTC | RT-qPCR analysis |
| DXR-F | TCCAGGAGAGCAAGGAGTGA | RT-qPCR analysis |
| DXR-R | TGCAGCAACCGTAGGCTTTA | RT-qPCR analysis |
| MCT-F | CGGTTGGAGCAGCTGTACT | RT-qPCR analysis |
| MCT-R | TCACCTGTGGTGTCTGCATT | RT-qPCR analysis |
| FPPS1-F | TAAACCCGACCCATCGAACG | RT-qPCR analysis |
| FPPS1-R | TTGGTGTCCCTCAATCGCTC | RT-qPCR analysis |
| FPPS2-F | TTGCTCATGGCGGGAGAAAA | RT-qPCR analysis |
| FPPS2-R | TGCCAAGTGTCTCAGGATCA | RT-qPCR analysis |
| AtCPT3_F | GCGCTTATGTCGATGCTG | RT-qPCR analysis |
| AtCPT3_R | CAGACTCAACCTCCTCAGG | RT-qPCR analysis |
| qPPR2F_2 | GTCACGCCATTCTTGGATCG | RT-qPCR analysis for PPRD2 |
| qPPR2R_2 | TCGCAATCAGTAAACCCGCA | RT-qPCR analysis for PPRD2 |
| PP2A-F | TAACGTGGCCAAAATGATGC | RT-qPCR analysis |
| PP2A-R | GTTCTCCACAACCGCTTGGT | RT-qPCR analysis |


Cho, Y., & Kanehara, K. (2017). Endoplasmic reticulum stress response in Arabidopsis roots. *Frontiers in Plant Science*, *8*, 225027. https://doi.org/10.3389/FPLS.2017.00144/BIBTEX

Jung, H., Lee, H. N., Marshall, R. S., Lomax, A. W., Yoon, M. J., Kim, J., Kim, J. H., Vierstra, R. D., & Chung, T. (2020). Arabidopsis cargo receptor NBR1 mediates selective autophagy of defective proteins. *Journal of Experimental Botany*, *71*(1), 73–89. https://doi.org/10.1093/JXB/ERZ404

Pastor-Cantizano, N., Bernat-Silvestre, C., Marcote, M. J., & Aniento, F. (2018). Loss of Arabidopsis p24 function affects ERD2 trafficking and Golgi structure, and activates the unfolded protein response. *Journal of Cell Science*, *131*(2). https://doi.org/10.1242/JCS.203802/258327/AM/LOSS-OF-ARABIDOPSIS-P24-FUNCTION-AFFECTS-ERD2
