## Supplemental Figures S1-S9 for "The *Arabidopsis thaliana* TRAPPIII subunit AtTRAPPC8/AtTRS85 is involved in ER functioning and autophagy"

by Hoffman-Sommer et al. (2024)

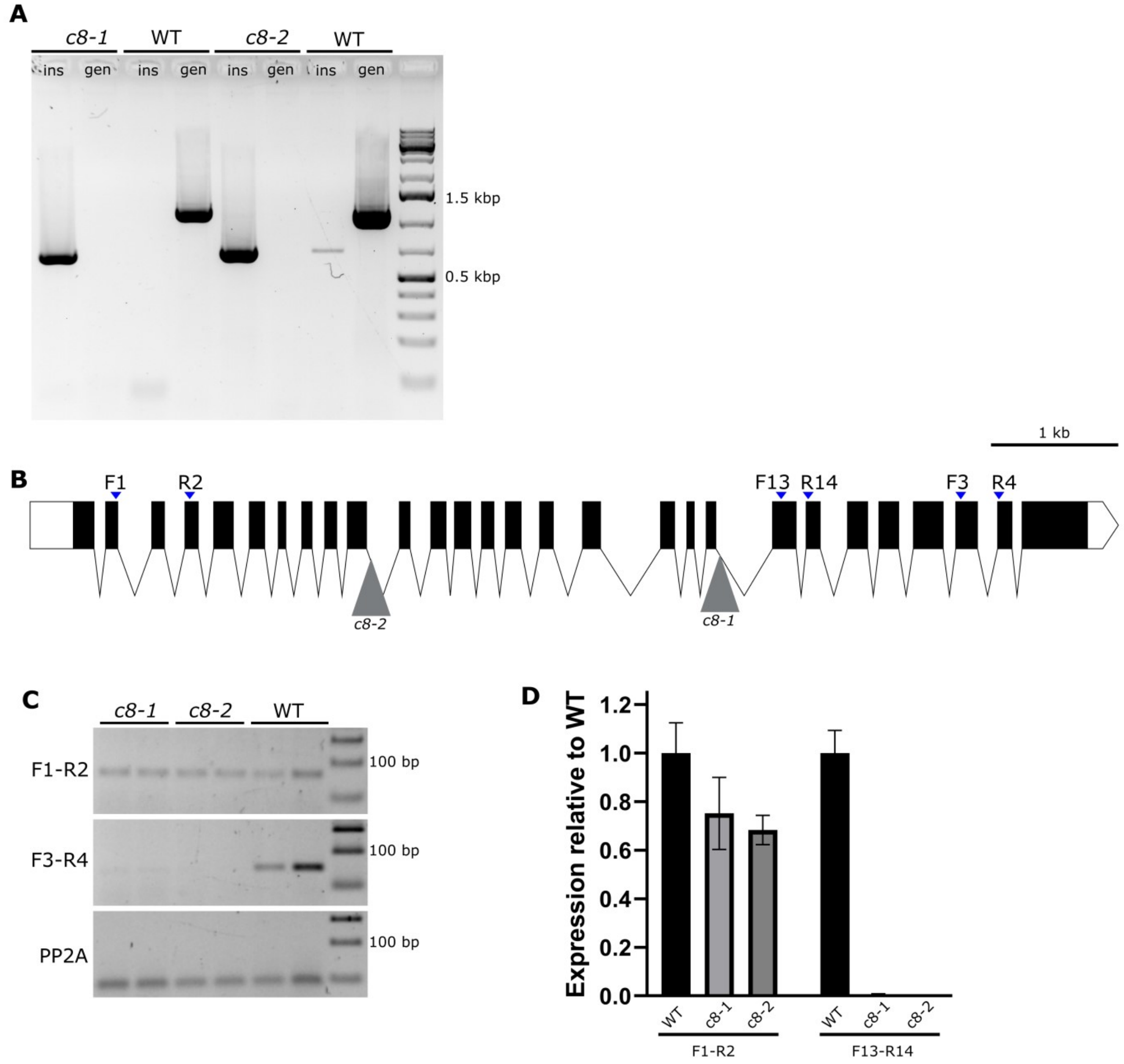


**Fig. S1.** Analysis of the insertion lines SALK_124093 (*trappc8-1*) and SALK_130580 (*trappc8-2*). (A) Genotyping of the isolated homozygotic mutant plants. ins – confirmation of allele with T-DNA insert, gen – confirmation of genomic allele. (B) Diagram of the intron-exon structure of the TRAPPC8 gene. The positioning of insertions in the mutant lines and of primers used for semi-quantitative PCR and RT-qPCR is shown. (C) Results of semi-quantitative PCR analyzing the expression of the mutated TRAPPC8 genes. PP2A served as an expression control. (D) RT-qPCR of the analyzed lines. The primer pairs used are indicated below the graph. RNA for experiments shown in C and D was extracted from rosette leaves of 4-week old plants grown in soil.

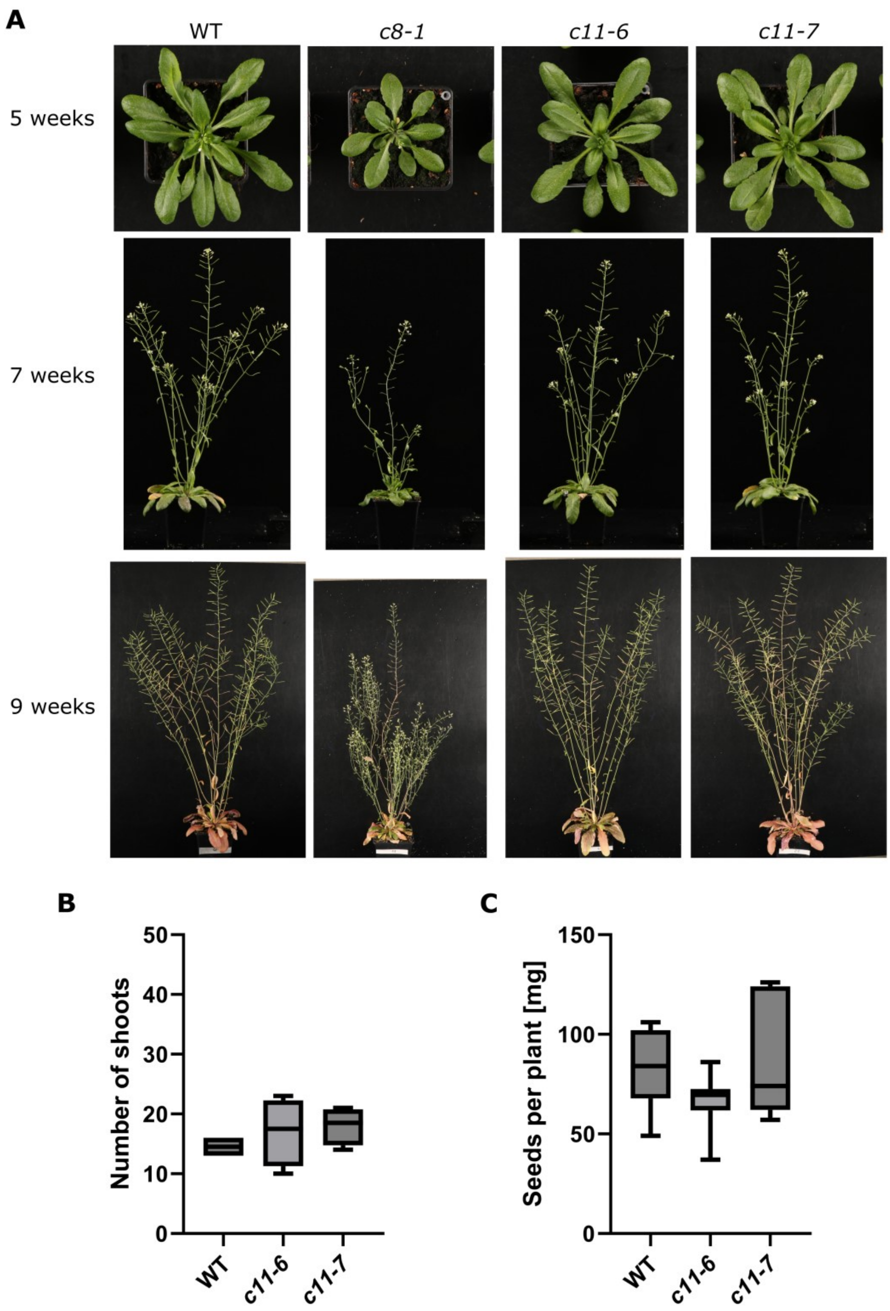


**Fig. S2.** Phenotypic analysis of the mutant lines *trappc11-6* and *trappc11-7*. (A) Overall morphology of WT and mutant plants cultured in soil. The mutant *trappc8-1* is shown for comparison. (B) Quantification of total number of shoots (primary and secondary) of 9-week old plants. 4 plants were scored for each genotype. (C) Quantification of amount of seeds produced per plant. The seeds of 7-8 individual plants were weighed for each genotype. The differences between WT and mutants were statistically insignificant in both B and C.

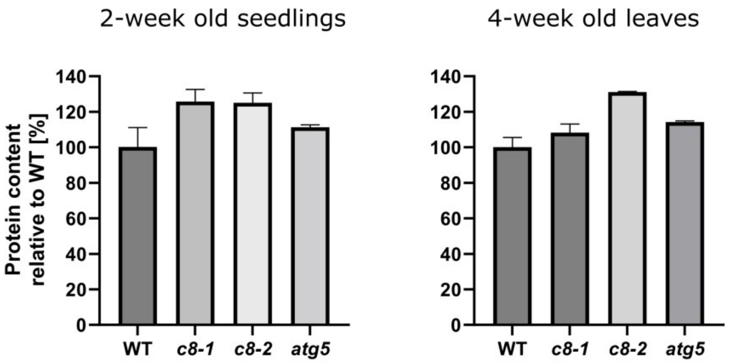


**Fig. S3.** Protein content per mg of tissue was assayed in 2-week old seedlings grown in liquid culture and in rosette leaves of 4-week old, soil-grown plants of the indicated genotypes. The differences between WT and mutant plants were not statistically significant for the control line *atg5*.

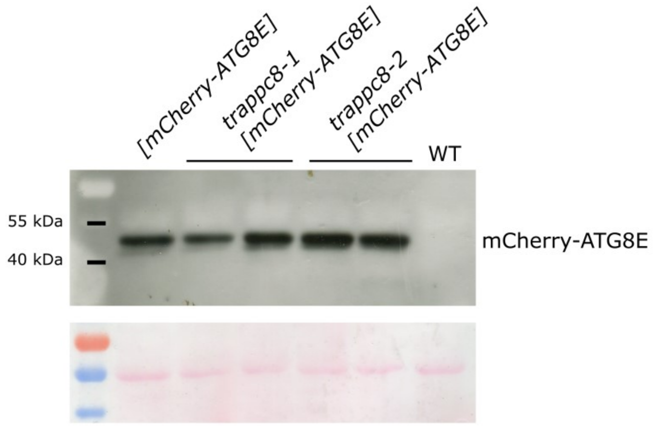


**Fig. S4.** Western blot confirming correct expression of the mCherry-ATG8E hybrid. Protein extracts were prepared from leaves of 5-week old plants grown in soil. The membrane was probed with an anti-mCherry antibody. The detected bands correspond to the expected molecular mass of mCherry-ATG8E, which is 46 kDa. Ponceau S staining of the membrane is shown below to demonstrate gel loading.

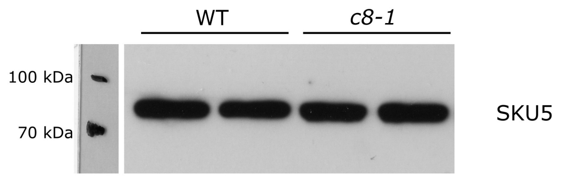


**Fig. S5.** Western blot with anti-SKU5 antibodies. Protein extracts were prepared from 2-week old seedlings grown in liquid culture.

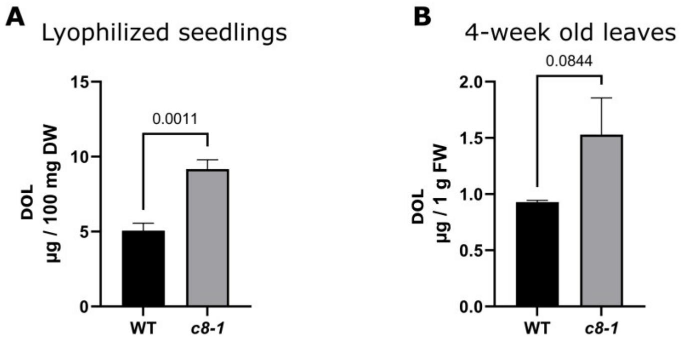


**Fig. S6.** Accumulation of dolichols (-15, -16 and -17) was assayed by HPLC-UV analysis of isoprenoids extracted from (A) lyophilized seedlings and (B) 4-week old leaves of WT plants and *trappc8-1* mutants. DW – dry weight, FW – fresh weight.

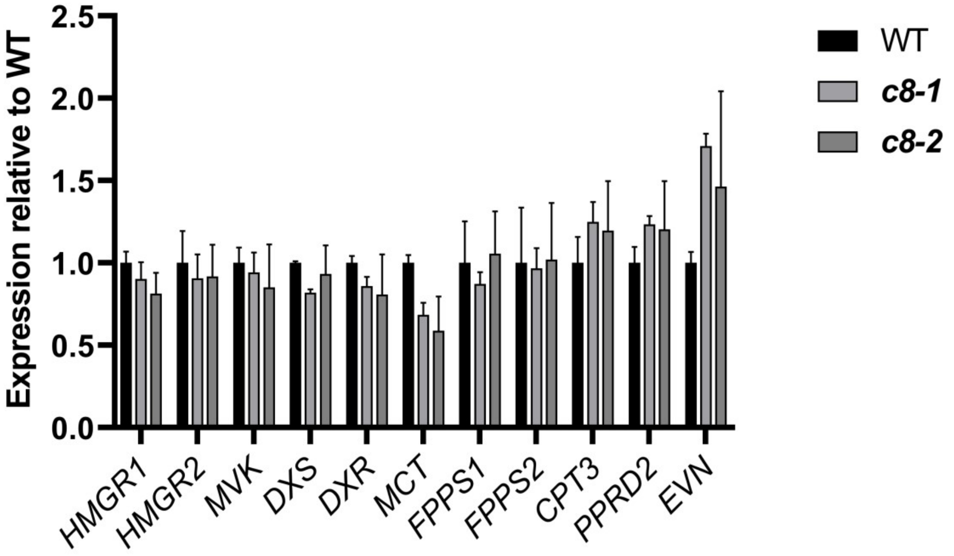


**Fig. S7.** Results of an RT-qPCR experiment analyzing the expression of genes from the dolichyl phosphate biosynthesis pathway in the mutants *trappc8-1* and *trappc8-2*. RNA for this experiment was prepared from 2-week old seedlings grown in liquid culture.

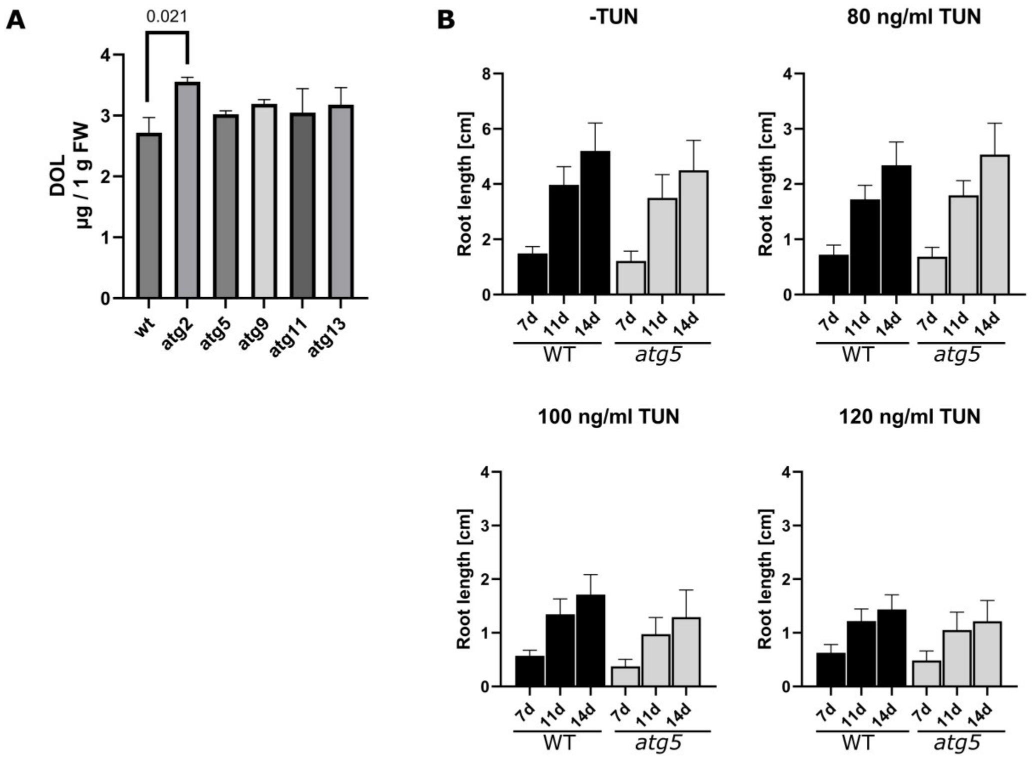


**Fig. S8.** Increased dolichol accumulation is not a feature of all autophagic mutants. (A) Dolichol levels were assayed by HPLC-UV analysis of isoprenoids extracted from 2-week old liquid-grown seedlings of the indicated genotypes. (B) WT and *atg5* seeds were sowed on plates containing the indicated concentrations of tunicamycin (TUN) in the growth medium. Seedlings were photographed after 7, 11 and 14 days of growth and root length was measured using ImageJ. No significant differences between WT and mutant seedlings were observed.

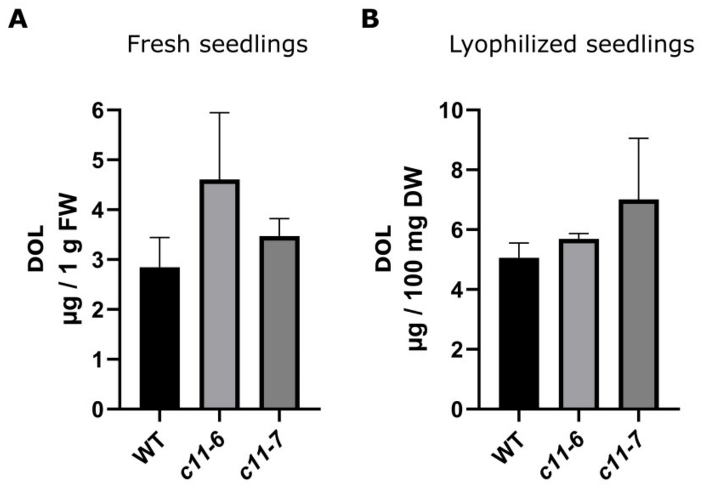


**Fig. S9.** Accumulation of dolichols (sum of dolichol-15, -16 and -17) was assayed by HPLC-UV analysis of isoprenoids extracted from seedlings of WT and *trappc11* lines cultured for 2 weeks in liquid medium and (A) used freshly or (B) lyophilized. FW – fresh weight, DW – dry weight. In this experiment all differences between lines were statistically not significant.
